# Genetic dissection of microglia cannibalism reveals an IL10 signaling axis controls microglia lifespan

**DOI:** 10.1101/2025.11.21.689277

**Authors:** Hannah Gordon, Dailin Gan, Antonio Dolojan, John Dennen, Camden A. Hoover, Zachary M. Koh, Khang Chau, Ricky Avalos Arceo, Cara Cavanaugh, Jun Li, Cody J. Smith

## Abstract

The development of complex organs, like the brain, demands a robust system for tissue remodeling and cellular debris clearance. In the brain, this function is performed by microglia, which must clear diverse debris substrates, including that caused by cell death. Although the subsequent fate of these phagocytic microglia is a critical regulatory point that impacts whether the brain resolves a debris environment, the genetic mechanisms that control microglia fate after debris clearance remain mostly unknown. To address this, we conducted a large-scale CRISPR screen in zebrafish using a custom-built robotic confocal microscope. We selected candidate genes from a single-cell RNA sequencing dataset of embryonic mouse microglia. This screen identified several modulators of microglial lifespan and cannibalism that are enriched in mouse and zebrafish microglia, including interleukin-10 receptor beta (*il10rb*), a receptor subunit for the cytokine IL10. Perturbation of *il10*, *il10rb,* and downstream signaling molecules JAK/STAT in zebrafish reduced microglial death. Expression analysis in mouse and zebrafish confirmed that microglia express both *il10* and *il10rb*. Given the established role of IL10 in lysosomal remodeling, we hypothesized that it regulates microglial survival through lysosomal acidification. While *il10rb* perturbation did not alter lysosome number or size, it caused a significant reduction in LysoTracker-positive lysosomes, indicating decreased lysosomal acidification. Inhibiting v-ATPase also reduced microglial death, reinforcing the link between lysosomal pH and cell fate. Our findings reveal a cytokine-regulated mechanism where lysosomal dynamics determine the survival of phagocytic microglia. We propose that a necroptosis-cannibalism process functions as a quality control mechanism for microglial turnover, which is critical for refining neuroimmune cell function in the brain.

## Introduction

The clearance of dead cells is a critical component of organ development and health. Microglia, the resident immune cell of the central nervous system (CNS)^1^, serve as the dedicated undertakers of the CNS, by continuously clearing the environment of cellular debris and dead cells^2,3^. One of their most fundamental roles during embryonic development is efferocytosis^4–11^, the swift and non-inflammatory clearance of dead and dying cells^12^. This clearance is crucial for maintaining brain homeostasis^8,13^ and tissue patterning^14^, as impaired efferocytosis by microglia can provoke inflammation^15,16^, lead to aberrant circuit function^17,18^, and drive neurological pathologies^11^. Despite this, the mechanisms regulating microglia themselves during periods of this intense phagocytic demand—and how their own survival is influenced by the debris they ingest—remain poorly understood.

While efferocytosis is typically a protective and homeostatic process^13,19^, the high-volume uptake of dead cells can impose significant metabolic^20–22^ and endolysosomal stress^23–25^ on phagocytes. This concept is supported by literature showing that macrophages and other phagocytic cells can undergo cell death due to debris overload in contexts ranging from atherosclerosis^26–28^ to spinal cord injury^29^. This death is not just a passive failure; it is likely an evolutionarily conserved mechanism to prevent prolonged cell damage or the propagation of cytotoxic proteins and undigested lipids within the tissue. In atherosclerosis, the continuous, high-volume uptake of oxidized lipids leads to the formation of lipid-laden (foamy) macrophages^30,31^. These phagocytically burdened macrophages are defective phagocytes^32,33^, and are prone to programmed cell death^34–36^, which represents a crucial, self-refining mechanism to eliminate overloaded cells and prevent the establishment of chronic, non-resolving inflammation^37^. This raises the intriguing possibility that their essential role in clearing debris comes at a cost to their own viability, and that the phagocytosis to cell death process is a self-limiting quality control mechanism. Indeed, recent work has revealed a paradoxical process where embryonic microglia die after becoming phagocytic, and their corpses are subsequently engulfed by other microglia in a process termed microglia cannibalism^38^. This form of microglial death appears to be mediated by necroptosis^38^, a regulated and inflammatory form of cell death characterized by plasma membrane rupture^39,40^. This previously unappreciated circuit of efferocytosis-induced necroptosis and cannibalism represents a key mechanism of microglial turnover and raises fundamental questions about its regulation and its developmental purpose.

To investigate how microglial lifespan and cannibalism are regulated under the high phagocytic load of neural development, we performed a high-throughput CRISPR-Cas9 screen in larval zebrafish using a dataset of embryonic mouse microglia^41^. This screen revealed that a loss of function in interleukin-10 receptor subunit beta (*il10rb*) significantly reduced cannibalism by decreasing microglial death. The cytokine interleukin-10 (IL10) is known for its potent immunosuppressive effects through the canonical JAK/STAT signaling pathway^42,43^, which is mediated by the accessory receptor subunit IL10RB that is expressed by microglia^41,44^. While generally associated with cell survival^45,46^, IL10 has also been shown to induce cell death under specific stress conditions^47–51^, presenting a potential mechanism of the role of IL10 in developmental turnover. Mechanistically, downstream signaling through STAT3 is linked to the endolysosomal system; STAT3 increases the activity of the vacuolar H^+^-ATPase (v-ATPase) pumps^52,53^ that are essential for maintaining the highly acidic environment of the lysosome^52,54,55^. Additionally, there is recent evidence linking IL10 and necroptosis. In a study looking at the effects of IL10 on neonatal mice with unilateral ureteral obstruction, a model for congenital obstructive nephropathies, *Il10^−/−^* mice showed a reduction in necroptosis^56^. However, whether IL10 signaling is the missing link that controls the necroptotic fate of cells after debris clearance remains unknown.

Here, we show that IL10 signaling is a central orchestrator of microglial turnover during development. We demonstrate that microglia express both *il10* and *il10rb*, and find that IL10 acts as a direct pro-death signal by engaging the IL10RB/JAK/STAT3 axis to drive necroptosis. Mechanistically, we show that disrupting this signaling impairs lysosomal acidification, a key step in debris degradation. This finding positions the lysosome as the critical stress checkpoint where high phagocytic load and IL10 signaling converge to determine cell fate. Furthermore, we demonstrate that exogenous IL10 accelerates microglial death. Perturbation of *il10rb* causes microglia to be longer-lived but also exhibit impaired debris clearance and pronounced morphological dysfunction following injury. These findings support a model in which IL10 signaling sensitizes phagocytic microglia to necroptosis by modulating lysosomal function, thereby linking immune regulation to a critical developmental cell death pathway that refines immune cell colonization in the developing brain.

## Results

Building on the recent finding that embryonic microglia undergo necroptosis following phagocytic engulfment of dead cells, we sought to identify the molecular regulators of microglial lifespan and their cannibalism. To identify the molecular components governing this essential turnover mechanism, we designed a large-scale CRISPR knockdown screen. Candidate genes were selected from a re-analysis of a single-cell RNA sequencing (scRNA-seq) dataset of E14.5 embryonic mouse brains^41^ (Fig. 1A). This analysis identified key clusters composed of microglia and their precursors (Fig. 1A). We focused on genes that were enriched in a cluster of cells that expressed transcripts consistent with immature microglia or their precursors and genes expressed in clusters of mature microglia (Fig. 1 A-B). These clusters expressed canonical microglial genes like *P2ry12* and *Tmem119* (Fig. 1C). Using differential enrichment of transcripts, we identified 556 transcripts in the mature microglia clusters and 137 enriched genes in the immature microglia cluster. To select genes for screening we prioritized first by identifying the transcripts most highly enriched throughout the dataset. Of those highly enriched transcripts, we then selected genes conserved in zebrafish. Finally, we honed primarily on genes coding for transcription factors (e.g. *Irf8, Maf, Nfe2l2, Pou2f2, Spi1*), cell-surface receptors (e.g. *Ccr1, Nrp1, P2rx4, F11r, Csf1r, Fcgr3*), and secreted proteins (e.g. *Gas6*, *Tgfb1, Ninj1, Mydgf*) due to their potential to drive large-scale changes in cell fate and communication. Since microglia are highly motile in order to scavenge and clear debris, we also included genes associated with cytoskeletal function to investigate their role in turnover (e.g. *Arpc2, Arpc1b, Capg, Pfn1*). The final screen list composed of 86 genes, 37 that were enriched only in the microglia precursor cluster, 43 that were enriched in mature microglia clusters, and 13 that were enriched in both mature and immature microglia (Table S1).

**Figure 1.**
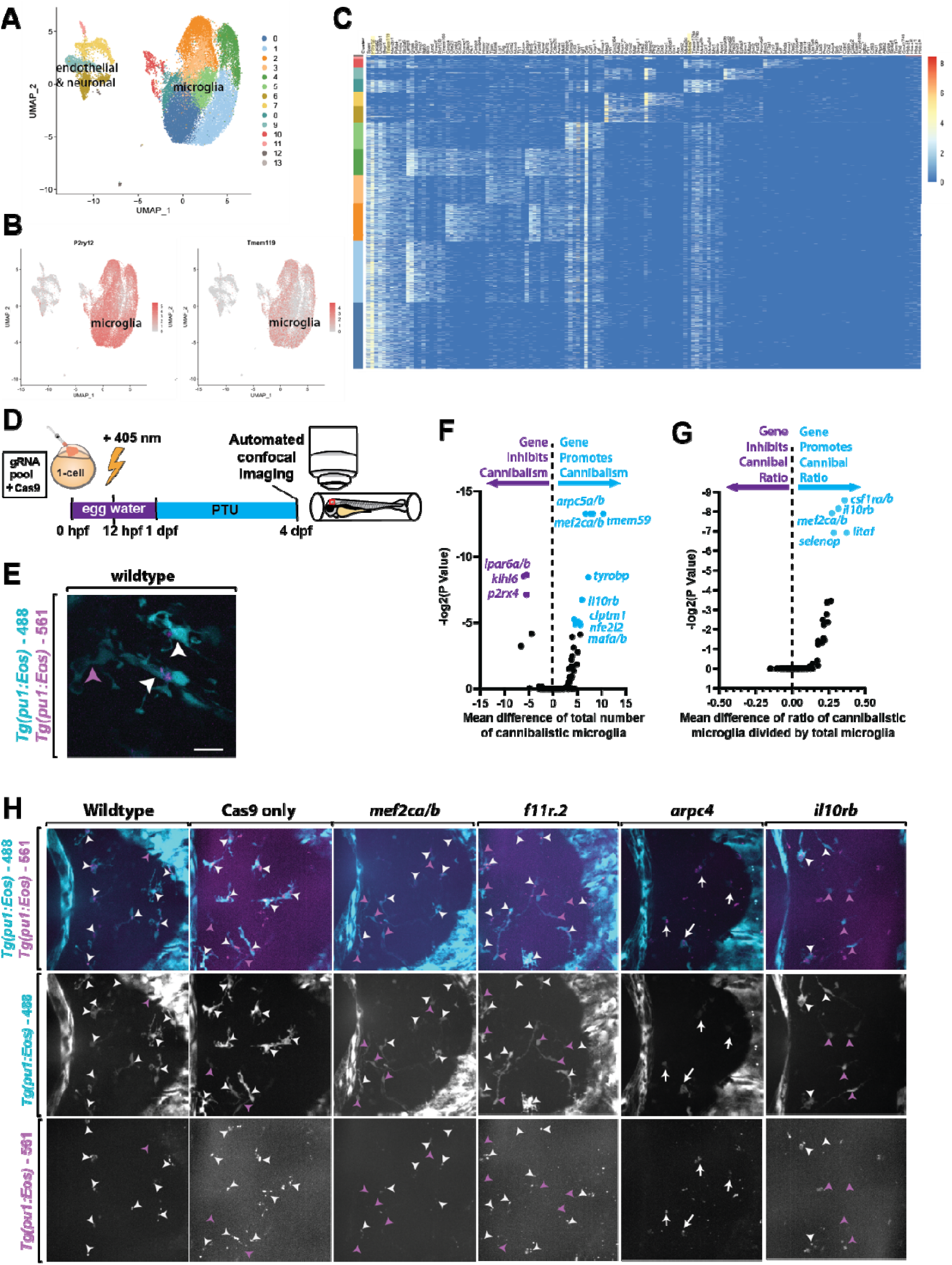
A CRISPR-Cas9 screen reveals novel regulators of microglial lifespan and self-clearance. (A) UMAP representing all 13 clusters identified in our re-analysis of the Hammond et al., 2019 sc-seq data of E14.5 mouse microglia. (B) Expression of canonical microglia genes *P2ry12* and *Tmem119* across all 13 clusters shown in the UMAP plot of mouse microglia. (C) Heat map representing expression levels across all clusters to identify distinct embryonic microglia clusters in mice at E14.5. (D) Schematic of experimental timeline for CRISPR knockdown screen. (E) Max z-projection of 5 dpf *Tg(pu1:Eos)* microglia showing cannibalistic microglia in zebrafish. White arrowheads represent debris containing cannibalistic microglia. Magenta arrowheads represent non-debris containing microglia. (F-G) Quantification of the mean difference of (F) the number of cannibalistic microglia and (G) the ratio of cannibalistic cells compared to the total microglia population in zebrafish. P value calculated with Two-way ANOVA post doc Tukey’s, Cas9 vs geneX gRNA. (H) Representative confocal z-projections of 5 dpf *Tg(pu1:Eos)* brains that were uninjected (wild-type), Cas9 only injected, or injected with Cas9 + gRNA for *mef2ca/b*, *f11r.2*, *arpc4*, or *il10rb*. Scale bar is 10 μm (E).

To quantitatively analyze microglial cannibalism, we used the transgenic zebrafish line *Tg*(*pu1:Eos*)^57^, which expresses the photoconvertible protein Eos under the control of the microglia/macrophage-specific *pu1* promoter. We achieved dual-color labeling of microglial progenitors by exposing zebrafish embryos to a 405 nm LED light array at 12 hours post-fertilization (hpf) (Fig. 1D) to selectively label a subset of microglia precursors that were expressing *pu1* at 12 hpf^58,59^. Since some microglia precursors initiate *pu1* expression after 12 hpf^60^, this early photoconversion created a mixed population of microglia at 4 dpf: some with only 488 Eos (these were non-photoconverted microglia that are pseudocolored cyan throughout images) and others with both 488 and 561 Eos (these were photoconverted microglia that are pseudocolored magenta throughout images). Embryos were grown to 4 dpf and imaged using a custom-built high-throughput robotic confocal microscope (Fig. 1D). The robot automatically positioned the midbrain of each animal to capture a 40 μm confocal z-stack.

To quantify the effect of gene knockdown on microglia cannibalism, we quantified each image for the number of 488-Eos/cyan microglia that either contained 561-Eos/magenta debris (indicating a cannibalistic microglia) or did not contain 561-Eos debris (indicating a non-cannibalistic microglia) (Fig. 1D). This allowed us to quantify the number of cannibalistic microglia and the ratio of cannibalistic cells to the total microglial population. Our initial screen identified 19 genes that, when knocked down, significantly altered the number of cannibalistic microglia and/or ratio of cannibalistic microglia compared to the total population of microglia (Fig. 1F-H, Fig. S1A-B). The dominant outcome of the screen was the identification of genes required for efficient cannibalism. Notably, very few genes were found to increase either the raw number of cannibalistic microglia or the ratio of cannibalistic microglia (Fig. 1F-H, Fig. S1A-B). However, the presence of genes that both increase and decrease the number of cannibalistic microglia could suggest a network of genes that balance microglia cannibalism. Collectively, the dual nature of these hits, with genes promoting and restricting cannibalism, suggests that microglia function is not controlled by a simple linear pathway, but rather by a complex regulatory network of genes that fine-tune microglial phagocytic activity during development.

To confirm the top hits from our initial CRISPR-Cas9 screen, we performed a secondary validation screen on 18 genes that showed a statistically significant phenotype (P<0.05). We excluded *csfr1a/b* from our validation screening, despite the significant decrease in the ratio of cannibalistic microglia that we saw (Fig. 1F), because of its known dominant role in microglia colonization^61^. We used the same Tg(*pu1:Eos*) line and high-throughput workflow (Fig. 1C) to quantify the number (Fig. 2A) and ratio (Fig. 2B) of cannibalistic microglia after knocking down each gene individually. The majority of the validated hits were found to promote cannibalism, as their perturbation resulted in a significant reduction in both the number and ratio of cannibalistic microglia (Fig. 2A-B). The validation screen successfully reproduced the phenotypes for 10 of the 18 genes, confirming their roles as robust regulators of microglial cannibalism.

**Figure 2.**
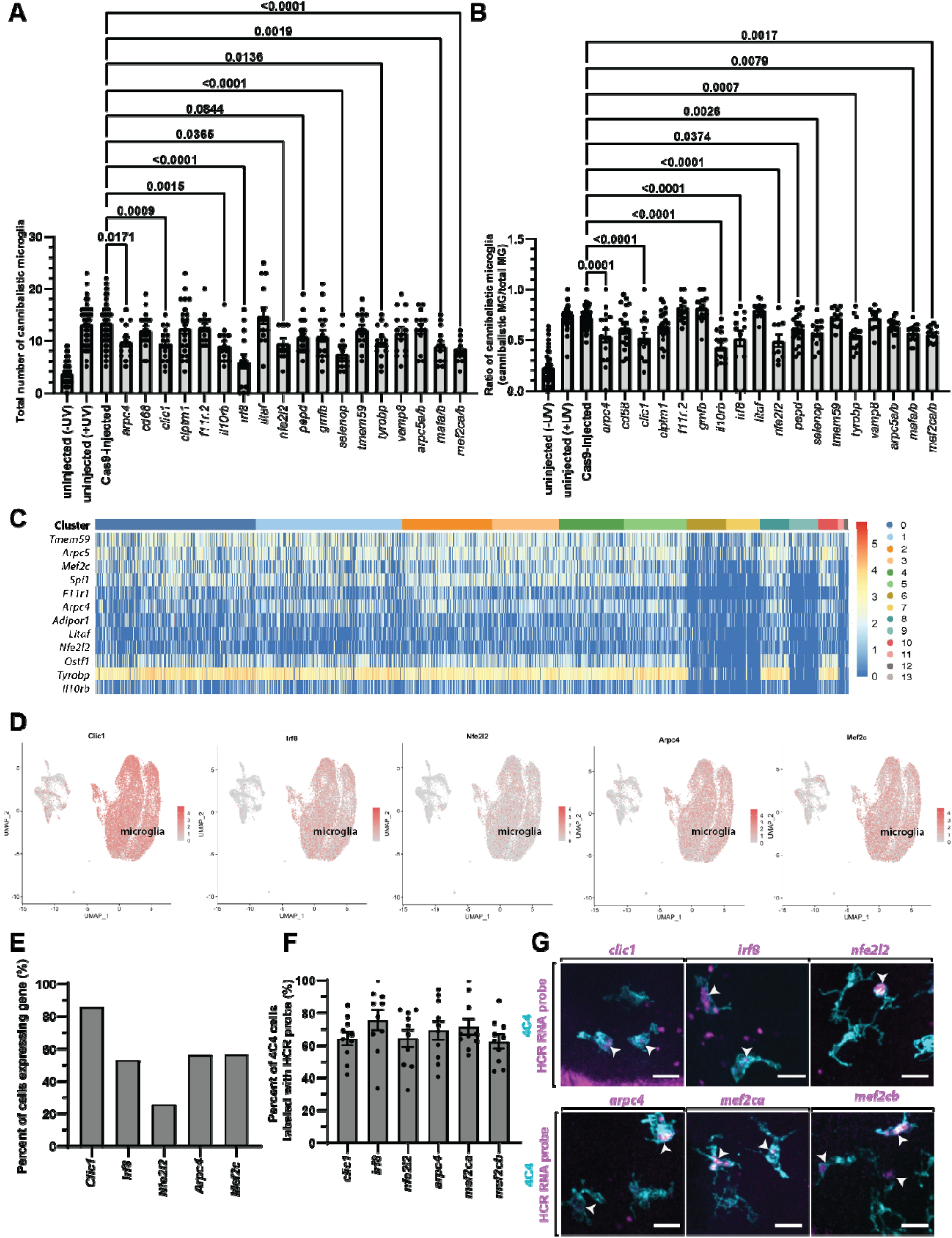
Validation and expression of candidate regulators of microglial cannibalism. (A-B) Quantification of (A) the total number of cannibalistic microglia (Cas9-injected vs. uninjected (−UV) p < 0.0001, Cas9-injected vs. uninjected (+UV) p > 0.9999, Cas9-injected vs. *arpc4* p = 0.0171, Cas9-injected vs. *cd68* p = 0.931, Cas9-injected vs. *clic1* p = 0.0009, Cas9-injected vs. *clptm1* p = 0.9951, Cas9-injected vs. *f11r.2* p > 0.9999, Cas9-injected vs. *gmfb* p = 0.1903, Cas9-injected vs. *il10rb* p = 0.0015, Cas9-injected vs. *irf8* p < 0.0001, Cas9-injected vs. *litaf* p = 0.9865, Cas9-injected vs. *nfe2l2* p = 0.0365, Cas9-injected vs. *pepd* p = 0.0844, Cas9-injected vs. *selenop* p < 0.0001, Cas9-injected vs. *tmem59* p = 0.9916, Cas9-injected vs. *tyrobp* p = 0.0136, Cas9-injected vs. *vamp8* p = 0.751, Cas9-injected vs. *arpc5a/b* p = 0.9956, Cas9-injected vs. *mafa/b* p = 0.0019, Dunnett’s multiple comparisons) and (B) the ratio of cannibalistic microglia (Cas9-injected vs. uninjected (−UV) p < 0.0001, Cas9-injected vs. uninjected (+UV) p > 0.9999, Cas9-injected vs. *arpc4* p = 0.0001, Cas9-injected vs. *cd68* p = 0.0781, Cas9-injected vs. *clic1* p < 0.0001, Cas9-injected vs. *clptm1* p = 0.1957, Cas9-injected vs. *f11r.2* p = 0.7573, Cas9-injected vs. *gmfb* p = 0.2435, Cas9-injected vs. *il10rb* p < 0.0001, Cas9-injected vs. *irf8* p < 0.0001, Cas9-injected vs. *litaf* p = 0.7613, Cas9-injected vs. *nfe2l2* p < 0.0001, Cas9-injected vs. *pepd* p = 0.0374, Cas9-injected vs. *selenop* p = 0.0026, Cas9-injected vs. *tmem59* p > 0.9999, Cas9-injected vs. *tyrobp* p = 0.0007, Cas9-injected vs. *vamp8* p > 0.9999, Cas9-injected vs. *arpc5a/b* p = 0.4278, Cas9-injected vs. *mafa/b* p = 0.0079, Cas9-injected vs. *mef2ca/b* p = 0.0017., Dunnett’s multiple comparisons) in 4 dpf *Tg(pu1:Eos)* knockdown animals and control animals. (C) Heatmap expression representing gene expression of validated knockdown genes among the 13 clusters. (D) UMAPs showing expression data in mouse microglia clusters of six validated knockdown genes (*Clic1, Irf8, Nfe2l2, Arpc4,* and *Mef2c*). (E) Quantification of the percent of cells in microglia clusters expressing *Clic1, Irf8, Nfe2l2, Arpc4,* and *Mef2c*. (F) Quantification of the percent of 4C4^+^ cells in zebrafish that are labeled with HCR RNA probes for *clic1, irf8, nfe2l2, arpc4, mef2ca,* and *mef2cb* (G) Representative confocal z-projections of 4 dpf animals stained for 4C4 and HCR RNA probes for *clic1, irf8, nfe2l2, arpc4, mef2ca* and *mef2cb*. White arrowheads represent microglia that express the HCR probe of interest. Scale bar represents 10 μm (G). Two-way ANOVA posthoc Dunnet’s multiple comparison (A,B).

Analysis of the validated genes revealed that microglial turnover is governed by a diverse, multiplex regulatory network rather than a single pathway. All listed genes were successfully validated as significant regulators of microglial cannibalism, and they represented a broad array of functional categories that highlight this complex control. This network includes master transcription factors vital for microglial identity and cell fate, such as *irf8*, *mafa/b*, and *mef2ca/b*, the latter being a noteworthy hit given its recently appreciated role in regulating microglial functions linked to autism risk and age-related disease^62^. Cytoskeletal genes like *arpc4*, which is responsible for actin polymerization, also significantly affected microglia cannibalism, likely due to defects in movement from cytoskeletal disorganization^63^. The validation of genes related to the phagocytosis/efferocytosis machinery, including *clic1*, which is known to be important for the acidification of phagolysosomes^64^, indicates that the physical process of engulfment and subsequent processing are tightly coupled. The identification of *f11r.2* (a junctional adhesion molecule) also underscores the role of membrane organization and adhesion in completing this process. Finally, the validation of genes coding for the transmembrane proteins and receptors *tyrobp* and *il10rb* reveal the crucial roles of cell-extrinsic signals in regulating the self-clearance process.

To determine if these genes are expressed in subtypes of embryonic microglia, we re-examined the scRNA-seq dataset^41^ of mouse microglia for five of the validated hits, including *clic1*, *irf8*, *nfe2l2*, *arpc4*, and *mef2ca/b*. Each of these genetic modifiers of microglia cannibalism were enriched throughout the mouse microglia transcriptional clusters. UMAPs of each genetic modifiers demonstrated their robust expression throughout the microglia clusters, albeit with different amounts for each modifier (Fig. 2C). A heatmap also shows gene expression of modifiers across all clusters to show specificity of expression in microglia (Fig. 2D). We also calculated the percentage of mouse microglia that expressed each genetic modifier (Fig. 2E), demonstrating that most mouse microglia at E14.5 express each genetic modifier. To determine if this gene expression was conserved in a system that we could thoroughly investigate microglia necroptosis and cannibalism, we tested expression *in vivo* using hybridization chain reaction RNA-FISH (HCR RNA-FISH) combined with immunohistochemistry (IHC) on 4 dpf zebrafish animals. To probe for expression in microglia, we stained for 4C4, a microglia specific antibody in zebrafish^65,66^, along with a set of probes for RNA expression of six genes that were hits in our screen. We generated HCR probes for *clic1*, *irf8*, *nfe2l2*, *arpc4*, and *mef2ca/b.* After staining, we imaged the midbrains of the 4 dpf animals and scored the percent of 4C4^+^ microglia that express each HCR probe per individual (Fig 2F-G). This revealed that all six genes were expressed in a majority of 4C4^+^ microglia in the brain, validating their conserved potential roles in microglial cannibalism and lifespan (Fig. 2F-G). Taken together, the conserved expression of these genetic modifiers in the majority of microglia across E14.5 mouse and 4 dpf zebrafish brains strongly validates their potential roles as core, conserved, regulators of microglial cannibalism and lifespan during development.

Among our validated hits, interleukin-10 receptor beta (*il10rb*) was a particularly compelling candidate. Its knockdown affected both the number of cannibalistic microglia and the ratio of cannibalistic microglia (Fig. 2A-B). Additionally, its canonical role in anti-inflammatory signaling and cell survival presented an intriguing paradox. Re-examining the scRNA-seq mouse dataset^41^, we found that *Il10rb* expression was highly specific to microglia clusters, with negligible expression in other brain cell types (Fig. 3A). We tested if this was conserved in zebrafish with HCR RNA-FISH and IHC, showing that approximately 87.88% ± 10.5% of 4C4-positive microglia expressed *il10rb* in 4 dpf animals (Fig. 3B-C). Given that IL10RB is a receptor for the cytokine interleukin-10 (IL10), we similarly explored the expression of *il10* in microglia to potentially suggest a cell-autonomous signaling cascade. HCR-FISH revealed that about 65.75% ± 15.46% of 4C4^+^ microglia expressed the ligand, *il10*, at 4 dpf (Fig. 3B-C). There were 4C4^-^ cells in the brain that also expressed *il10rb* and *il10* (Fig. 3C). This expression supports the hypothesis that *il10rb* is expressed by most developmental microglia and that IL10 signaling components are in the brain to potentially regulate microglia.

**Figure 3.**
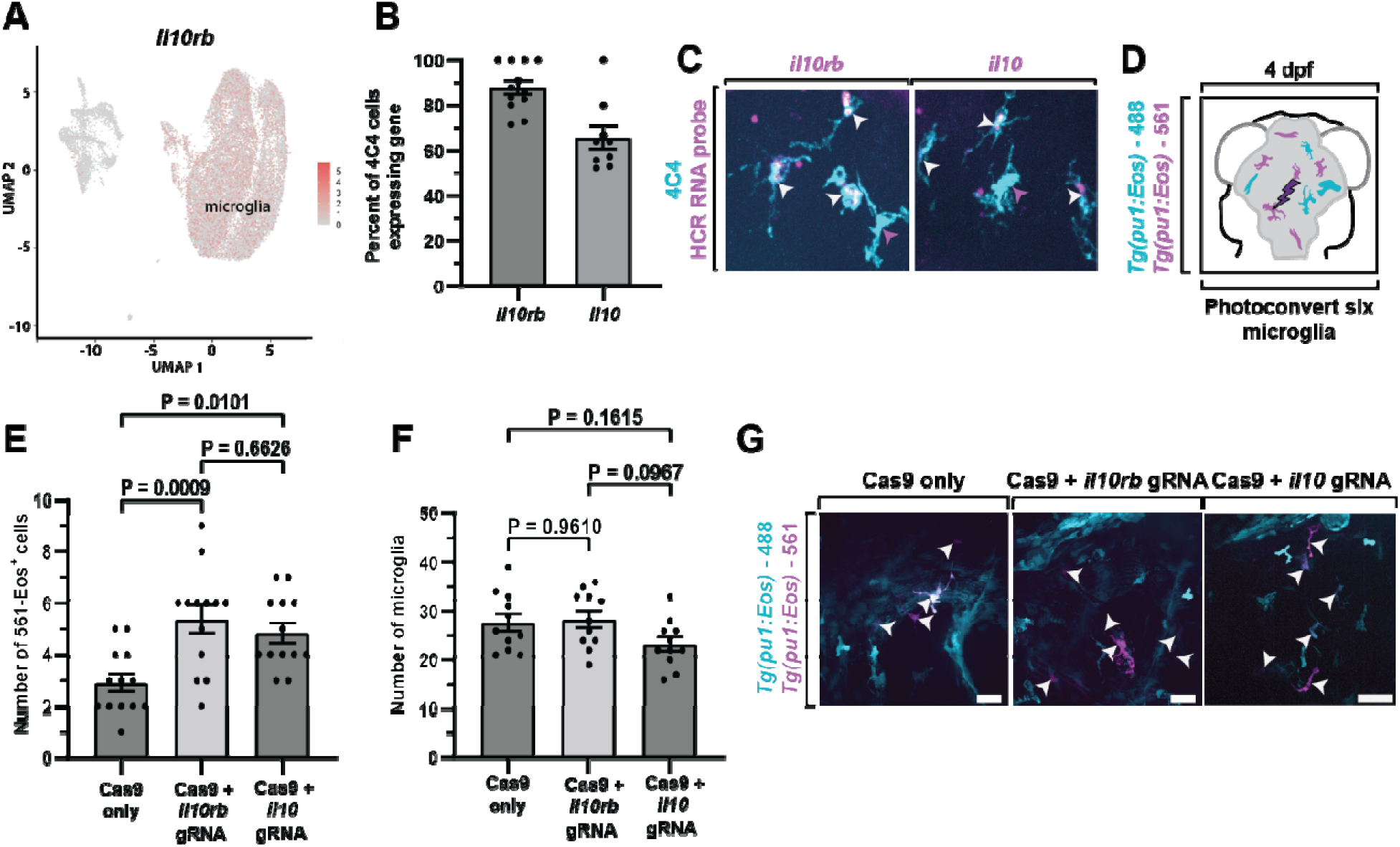
*il10rb* is a regulator of microglial lifespan. (A) Expression of *Il10rb* across all 13 clusters shown in the UMAP plot of mouse microglia. (B) Quantification of the percent of 4C4^+^ cells (microglia) expressing *il10rb* and *il10* in the zebrafish midbrain at 4 dpf. (C) Representative confocal z-projections showing 4C4^+^ microglia that express *il10rb* or *il10*. White arrowheads represent microglia that express the HCR probe of interest. Magenta arrowheads represent microglia that do not express the HCR probe of interest. (D) Schematic of experimental timeline for studying effect *il10rb* or *il10* knockdown on microglia lifespan and cannibalism. Six cells were targeted and photoconverted in 4 dpf *Tg(pu1:Eos)* animals that were injected with Cas9 only, Cas9 + *il10rb* gRNA, or Cas9 + *il10* gRNA. Animals were grown to 5 dpf and their brains were reimaged for the presence of photoconverted microglia and cannibal microglia. (E-F) Quantification of the number of (E) 561-Eos microglia (Cas9 only vs. Cas9 + *il10rb* gRNA p = 0.0009, Cas9 only vs. Cas9 + *il10* gRNA p = 0.0101, Cas9 + *il10rb* gRNA vs. Cas9 + *il10* gRNA p = 0.6626, Tukey’s multiple comparisons test) and (F) total microglia (Cas9 only vs. Cas9 + *il10rb* gRNA p = 0.961, Cas9 only vs. Cas9 + *il10* gRNA p = 0.1615, Cas9 + *il10rb* gRNA vs. Cas9 + *il10* gRNA p = 0.0967, Tukey’s multiple comparisons test) in 5 dpf animals injected with Cas9 only, Cas9 + *il10rb* gRNA, or Cas9 + *il10* gRNA. (G) Representative confocal z-projections of 5 dpf *Tg(pu1:Eos)* animals injected Cas9 only, Cas9 + *il10rb* gRNA, or Cas9 + *il10* gRNA. White arrowheads represent 561-Eos+/photoconverted microglia. Scale bars represent 20 μm (G).

While our screen quantified cannibalism, this metric serves as an indirect readout of microglial turnover. A lower amount of cannibalistic microglia could indicate two most likely yet distinct biological possibilities: 1) Microglial death was reduced, meaning fewer corpses were available for clearance, or 2) The efficiency of efferocytosis (cannibalism) was impaired, meaning corpses were present but were not cleared. To distinguish between these possibilities for *il10rb*, we performed a survival assay after perturbation of *il10* and *il10rb* with CRISPR. Guide efficiency for the CRISPR perturbation was confirmed by T7 endonuclease assay, which had an 87% indel efficiency (Fig. S2A). Only animals positive for an indel were used for quantification. We used *Tg(pu1:Eos)* animals and photoconverted six midbrain microglia at 4 dpf. We then re-imaged the same animals at 5 dpf to track the survival of the labeled cells (Fig. 3D). Control animals injected with Cas9 only animals retained an average of 2.9 ± 1.3 561-Eos cells, consistent with ongoing developmental cell death^38^. In contrast, Cas9 plus *il10rb* or *il10* gRNA animals retained significantly more 561-Eos cells (5.4 ± 2.0 and 4.8 ± 1.4, respectively) (Fig. 3E). While it is possible that this increased cell persistence was associated with changes in the total microglial population at 5 dpf, scoring the total microglia abundance in each perturbation showed the statistically indistinguishable abundance of microglia, indicating that the effect was specific to individual cell lifespan rather than colonization or proliferation (Fig. 3F-G). Additionally, when the number of cannibalistic microglia were quantified at 5 dpf, there were fewer cannibalistic microglia in Cas9 plus *il10*rb or *il10* gRNA injected animals compared to Cas9 only injected animals (Fig. S2B) (4.5 ± 1.9 cannibalistic microglia in Cas9 only; 1.6 ± 1.3 cannibalistic microglia in Cas9 + *il10rb* gRNA; 2.2 ± 1.8 cannibalistic microglia in Cas9 + *il10* gRNA). These results decisively separate the two possibilities, indicating that *il10* and its receptor *il10rb* are necessary for developmental microglial death and that the reduction in cannibalism observed in our screen is a secondary consequence of the microglia surviving longer, suggesting that IL10 acts as a critical regulator of microglial turnover rate during development.

Given that perturbing *il10rb* or *il10* increased the number of live, 561-Eos microglia at 5 dpf, we next investigated whether IL10 signaling actively promotes the developmental death of microglia. To directly test if exogenous IL10 could drive microglial death *in vivo*, we photoconverted six microglia in the midbrain of 4 dpf *Tg(pu1:Eos)* animals and immediately injected either recombinant human IL10 (rhIL10) or PBS (vehicle control) into the brain ventricles (Fig. 4A) We utilized recombinant human protein to further test the potential conservation of this process. We re-imaged the animals at 5 dpf to assess the fate of the labeled cells. In PBS-injected controls, an average of 4.0 ± 0.8 561-Eos microglia persisted, while rhIL10-injected animals retained only 2.1 ± 1.2 561-Eos microglia (Fig. 4B). This indicates that IL10 exposure significantly accelerated microglial removal between 4 and 5 dpf. To further confirm cell death, we quantified the number of cannibal microglia containing the remnants of dead microglia. rhIL10 injected animals had an average of 6.4 ± 2.8 cannibal microglia, double the 3.1 ± 1.1 cannibal microglia observed in PBS controls (Fig. 4C-D). Notably, total microglial abundance was unchanged, indicating that rhIL10 caused a selective loss of the labeled population rather than affecting colonization or proliferation (Fig. S2C).

**Figure 4.**
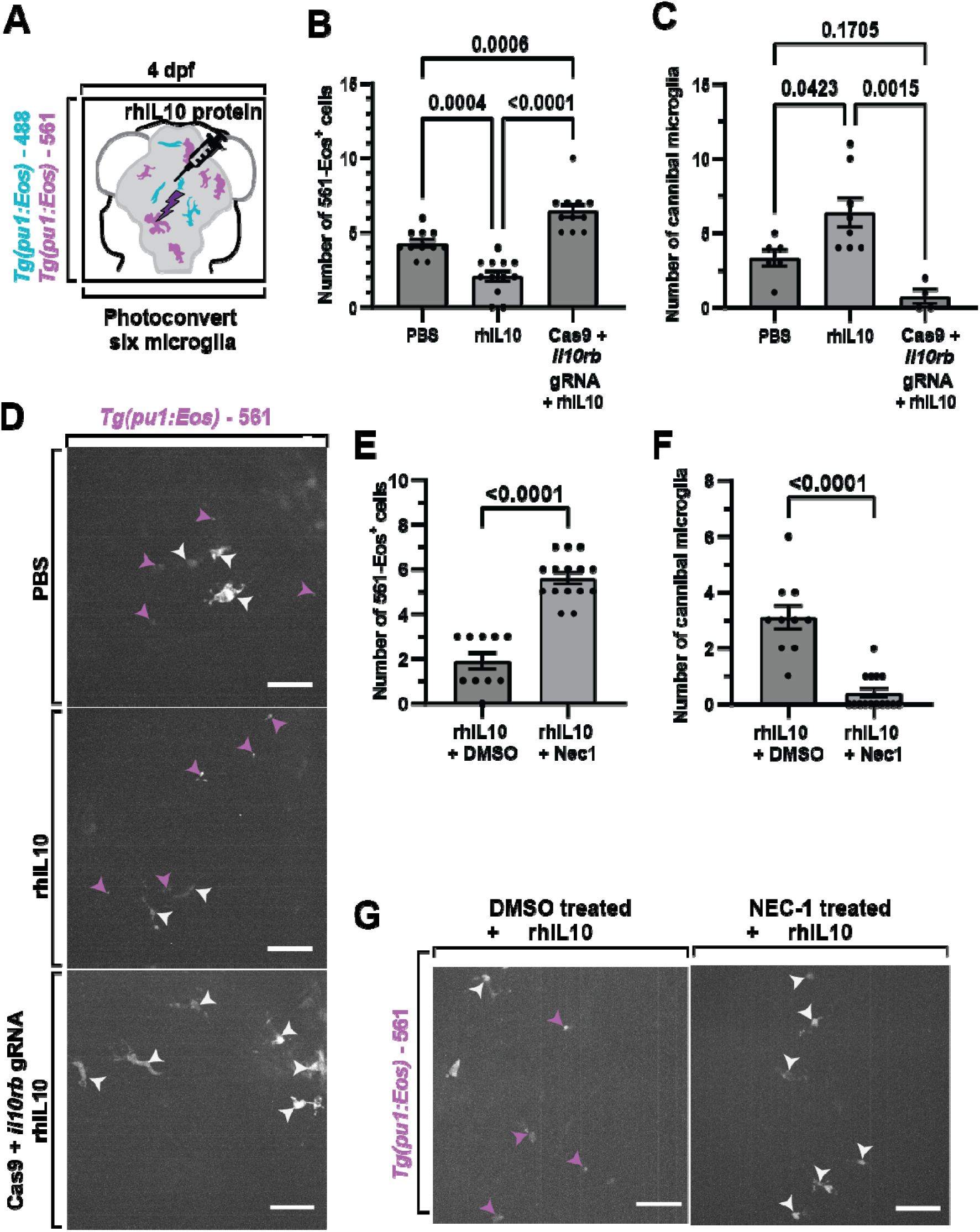
IL10 signaling drives microglial death via necroptosis. (A) Schematic of experimental timeline for studying effect of increased IL-10 signaling on microglia lifespan and cannibalism. Six cells were targeted and photoconverted in 4 dpf *Tg(pu1:Eos)* animals. After photoconversion, animals were injected with PBS or rhIL10 in their brain ventricles. Animals were grown to 5 dpf and their brains were reimaged for the presence of photoconverted microglia and cannibal microglia. (B-C) Quantification of the (B) number of 561-Eos microglia (PBS vs. rhIL10 p = 0.0004, PBS vs. Cas9 + *il10rb* gRNA + rhIL10 p = 0.0006, rhIL10 vs. Cas9 + *il10rb* gRNA + rhIL10 p < 0.0001; Tukey’s multiple comparisons test) and (C) number of cannibalistic microglia (PBS vs. rhIL10 p = 0.0423, PBS vs. Cas9 + *il10rb* gRNA + rhIL10 p = 0.1705, rhIL10 vs. Cas9 + *il10rb* gRNA + rhIL10 p = 0.0015; Tukey’s multiple comparisons test) in 5 dpf animals injected with PBS or rhIL10. (D) Representative confocal z-projections of 5 dpf *Tg(pu1:Eos)* animals injected with PBS or rhIL10 showing photoconverted microglia and the presence of cannibal microglia. (E-F) Quantification of the (E) number of 561-Eos (rhIL10 +DMSO vs rhIL10 + Nec1 p < 0.0001; unpaired t-test) and (F) number of cannibalistic microglia (rhIL10 +DMSO vs rhIL10 + Nec1 p < 0.0001; unpaired t-test) in 5 dpf animals injected with rhIL10 and bath treated with DMSO or Nec1. (G) Representative confocal z-projections showing 5 dpf *Tg(pu1:Eos)* animals injected with rhIL10 and then bath treated with DMSO or Nec1 showing photoconverted microglia and photoconverted debris/presence of cannibalistic microglia. White arrowheads denote 561-Eos^+^ photoconverted microglia (D,G). Magenta arrowheads denote 561-Eos^+^ debris containing cannibalistic microglia (D,G). Scale bars represent 20 μm (D,G).

If IL10 functions through the IL10RB receptor, we would expect that the effect of IL10 on microglia death would be eliminated in *il10rb* knockdown. To test this, we repeated the rhIL10 injection in Cas9 plus *il10rb* gRNA injected animals. As expected, the loss of receptor function abolished the pro-death effect. rhIL10 injection failed to reduce the number of 561-Eos microglia in *il10rb* gRNA injected animals (average 6.5 ± 1.4 561-Eos microglia), and yielded almost no cannibal microglia (0.4 ± 0.6 cannibalistic microglia) (Fig. 4 B-D). These results confirm that IL10 is sufficient to promote microglial death *in vivo*, and this activity is dependent on IL10RB-mediated signaling.

With the role of IL10 in microglia death established, we next sought to identify the specific mechanism of cell death caused by IL10 signaling. We first tested for apoptosis, a common form of programmed cell death, by performing immunohistochemistry for the active form of caspase-3, cleaved caspase-3 (cCASP3), a canonical apoptotic marker. We found no cCASP3^+^ microglia in either PBS- or rhIL10-injected animals (Fig. S2D), indicating that apoptosis is not the primary mode of death. To confirm the effect was specific to microglia and not to other neural cell types, we also quantified the total number of cCASP3^+^ cells in the brain and found no change (34.4 ± 7.6 cCASP3^+^ cells per PBS injected animal; 38.9 ± 8.4 cCASP3^+^ cells per rhIL10 injected animal), suggesting IL10 is not acting on other neural cell types through caspase-mediation (Fig. S2D-E).

We previously discovered that debris-burdened microglia die via necroptosis, and their corpses are subsequently cleared by neighboring microglia in a process termed microglia cannibalism^38^. Given that embryonic microglia are known to undergo necroptosis, we hypothesized that IL10 might instead be acting through this pathway. To test this, we co-treated rhIL10-injected animals with a necrostatin-1 (Nec-1) bath treatment, a specific RIPK1 inhibitor that blocks necroptosis^67^. Nec-1 treatment successfully rescued microglial survival, with animals retaining an average of 5.6 ± 1.0 561-Eos microglia compared to DMSO treated animals that only retained an average of 1.9 ± 1.2 561-Eos microglia (Fig 4E). Additionally, the Nec-1 treated animals only had an average of 0.4 ± 0.6 cannibalistic microglia compared to DMSO treated animals that had an average of 3.1 ± 1.4 cannibalistic microglia (Fig 4F-G). These findings strongly suggest that IL10 promotes microglial death via a necroptotic pathway in an *il10rb*-dependent manner.

We next sought to determine the intracellular signaling cascade downstream of the IL10 receptor that leads to microglial death. Interleukin-10 receptor signaling is canonically known to activate the Janus kinase (JAK) and Signal Transducer and Activator of Transcription 3 (STAT3) pathway^43^. To test if this canonical pathway mediates the IL10-induced microglial death, we used a pharmacological approach to inhibit JAK and STAT3 activity.

We first photoconverted six microglia in the midbrain of 4 dpf *Tg(pu1:Eos)* animals. Immediately following photoconversion, animals were treated with either DMSO (vehicle control), a pan JAK inhibitor (JAKi), or a STAT3 inhibitor (STAT3i). We then re-imaged the same animals at 5 dpf to assess the survival of the labeled cells (Fig 5A). In DMSO-treated control animals, we observed an average of 3.5 ± 1.3 561-Eos microglia remaining, consistent with the ongoing cell death seen in previous experiments (Fig. 5B, Fig. 3E). In stark contrast, animals treated with the STAT3i or JAKi retained an average of 5.8 ± 1.1 and 5.7 ± 0.8 561-Eos cells, respectively (Fig 5B). These results strongly suggest that both JAK and STAT3 signaling are required for the natural, developmental loss of microglia. Additionally, we quantified the number of cannibalistic microglia. JAKi and STAT3i treatment resulted in almost no cannibalistic microglia (an average of 0.2 ± 0.4 and 0.1 ± 0.3 cannibalistic microglia per animal, respectively), while DMSO-treated controls had an average of 2.4 ± 1.4 cannibalistic cells per animal, further supporting the conclusion that inhibiting this pathway prevents microglial death (Fig 5C-D).

**Figure 5.**
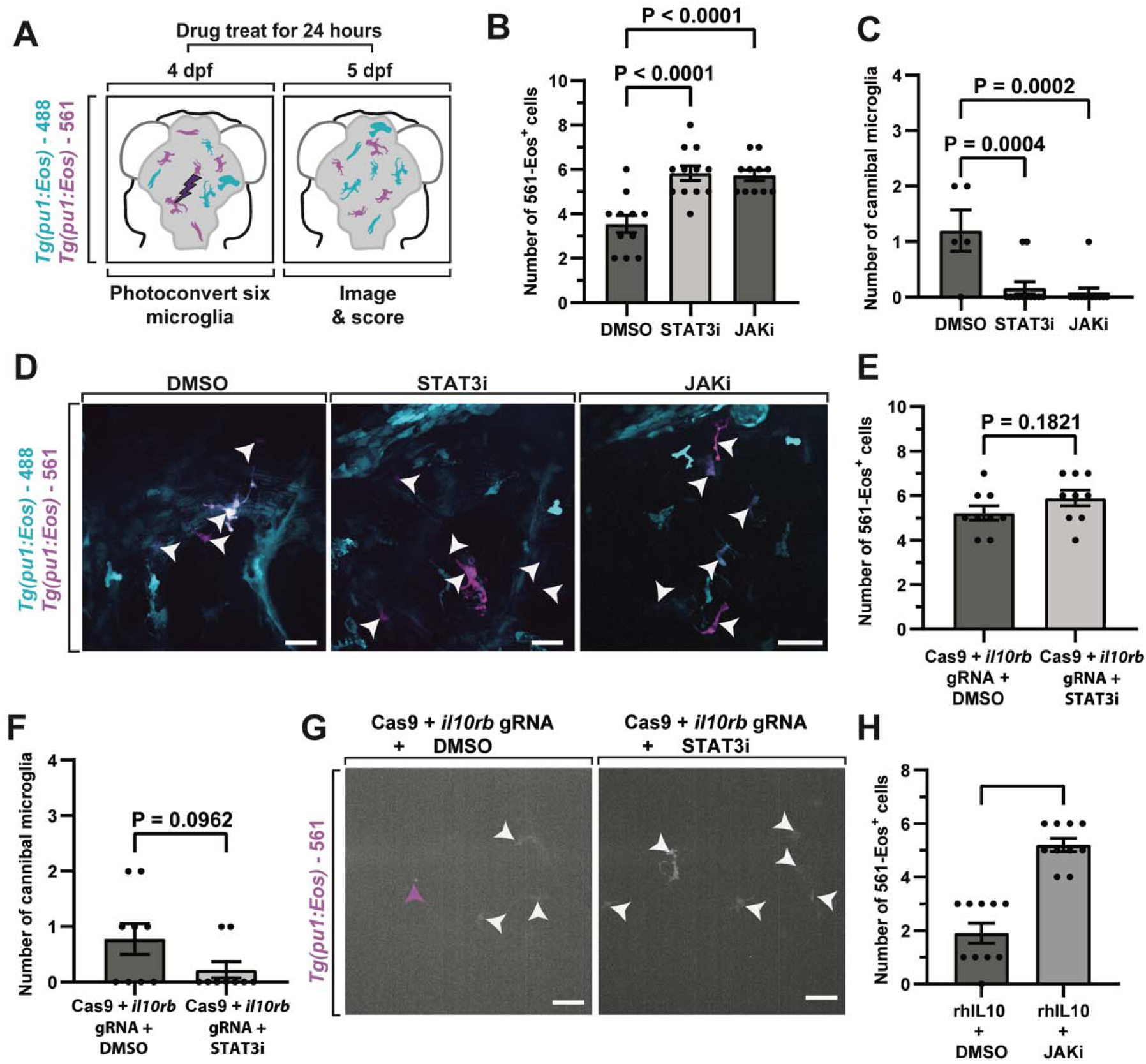
IL10 Signaling Promotes Microglial Death Through the JAK/STAT Pathway. (A) Schematic of experimental timeline for studying effect of blocking JAK/STAT signaling on microglia lifespan and cannibalism. Six cells were targeted and photoconverted in 4 dpf *Tg(pu1:Eos)* animals. After photoconversion, animals were bath treated with DMSO, STAT3i, or JAKi. Animals were grown to 5 dpf and their brains were reimaged for the presence of photoconverted microglia and cannibal microglia. (B-C) Quantification of the (B) number of 561-Eos microglia (DMSO vs STAT3i p < 0.0001, DMSO vs JAKi p < 0.0001; Dunnett’s multiple comparisons test) and (C) number of cannibal microglia (DMSO vs STAT3i p < 0.0001, DMSO vs JAKi p < 0.0001; Dunnett’s multiple comparisons test) in 5 dpf animals bath treated with DMSO, STAT3i, or JAKi. (D) Representative confocal z-projections of 5 dpf *Tg(pu1:Eos)* animals bath treated with DMSO, STAT3i, or JAKi showing photoconverted microglia and photoconverted debris containing presence of cannibalistic microglia. (E-F) Quantification of the (E) number of 561-Eos (cas9 + *il10rb* gRNA + DMSO vs cas9 + *il10rb* gRNA + STAT3i p = 0.1821; unpaired t-test) and (F) number of cannibal microglia (cas9 + *il10rb* gRNA + DMSO vs cas9 + *il10rb* gRNA + STAT3i p = 0.0962; unpaired t-test) in 5 dpf cas9 + *il10rb* gRNA bath treated with DMSO or STAT3i. (G) Representative confocal z-projections of 5 dpf *Tg(pu1:Eos)* cas9 + *il10rb* gRNA animals bath treated with DMSO or STAT3i showing photoconverted microglia and presence of cannibal microglia. (H) Quantification of the number of 561-Eos microglia in 5 dpf rhIL10 injected animals bath treated with DMSO or JAKi. White arrowheads denote 561-Eos microglia (D,G). Magenta arrowheads denote 561-Eos+ debris containing cannibal microglia (D,G). Scale bars represent 20 μm (D,G).

To confirm that JAK and STAT3 act in the same pathway as IL10RB, we first repeated the experiment in Cas9 plus *il10rb* gRNA injected animals. We treated Cas9 plus *il10rb* gRNA injected animals with either DMSO or STAT3i after photoconversion, and found no significant difference in the number of surviving 561-Eos microglia between the two groups (5.2 ± 1.0 561-Eos microglia in Cas9 plus *il10rb* gRNA injected + DMSO treated animals; 5.9 ± 1.0 561-Eos microglia in Cas9 plus *il10rb* gRNA injected + STAT3i treated animals) (Fig 5E). We also quantified the number of cannibalistic microglia and found no discernable difference between groups (0.8 ± 0.8 cannibalistic microglia Cas9 plus *il10rb* gRNA injected + DMSO treated animals; 0.2 ± 0.4 cannibalistic microglia Cas9 plus *il10rb* gRNA injected + STAT3i treated animals) (Fig 5F-G). The lack of an additive effect in the *il10rb* gRNA is consistent with the hypothesis that JAK activity is epistatic to IL10RB, supporting the idea that they function in the same signaling cascade.

Finally, to directly link IL10 signaling to JAK activity, we tested whether JAK inhibition could block the pro-death effect of exogenous IL10. We injected rhIL10 into the brain ventricles of 4 dpf animals and then bathed the animals in DMSO or JAKi water. We found that JAKi treatment successfully rescued microglial survival, with animals retaining a high number of 561-Eos microglia, a result similar to JAKi treatment alone (1.9 ± 1.2 561-Eos microglia in rhIL10 + DMSO treated animals; 5.2 ± 0.7 561-Eos microglia in rhIL10 + JAKi treated animals) (Fig 5H). This confirms that the pro-death effect of IL10 is mediated through a JAK-dependent signaling pathway. Collectively, these findings demonstrate that IL10 signaling promotes microglial necroptosis by activating the canonical JAK/STAT pathway. This establishes a clear molecular cascade, linking the extracellular cytokine signal to the intracellular machinery that governs microglial lifespan during development.

Our central model proposes that IL10 signaling acts as a quality control checkpoint, linking the phagocytic burden within a cell to the initiation of cell death. Having established that *il10rb* perturbation significantly reduces microglia death, we next investigated the relationship between IL10 signaling and the act of phagocytosis itself. Specifically, we tested if *il10rb*-mediated necroptosis required the presence of phagocytic activity. To address this, we performed a microglial survival assay on *il10rb* gRNA, treating them with either water (H_2_O) as a control or with O-phospho-L-serine (L-SOP), a known inhibitor of phagocytosis. L-SOP has been shown in our previous study to decrease microglia death^38^. Six microglia were photoconverted prior to drug treatment and animals were bath treated for 24 hours as done before (Fig. 5A). After 24 hours of treatment of L-SOP or H_2_O, we quantified the number of surviving photoconverted microglia and cannibal microglia in both groups. We found no significant difference in either the number of surviving 561-Eos cells (5.9 ± 1.1 561-Eos microglia per H_2_O treated *il10rb* gRNA; 5.8 ± 0.7 561-Eos microglia per L-SOP treated *il10rb* gRNA) or the number of cannibal microglia between the H2O- and L-SOP-treated *il10rb* gRNA (0.6 ± 0.7 cannibalistic microglia per H_2_O treated *il10rb* gRNA; 0.3 ± 0.5 cannibalistic microglia per L-SOP treated *il10rb* gRNA) (Fig 6A-C). This lack of an additive effect suggests that IL10 signaling acts downstream of, or in parallel with, the initial phagocytic engulfment signal, and that IL10RB and phagocytosis are components of the same cell biological process that regulates the final cell death decision.

**Figure 6.**
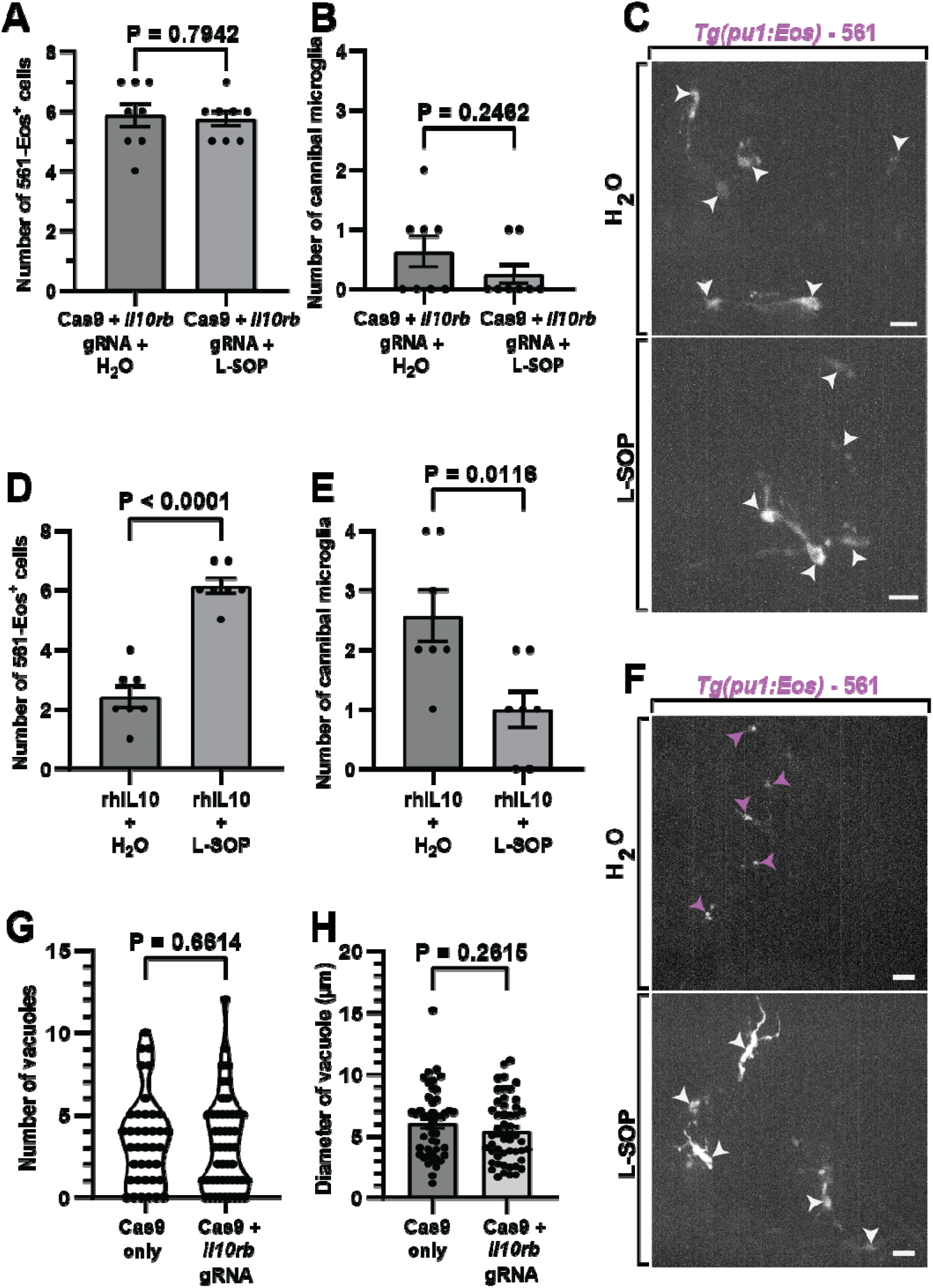
IL10 Signaling and Phagocytic Resolution. (A-B) Quantification of the (A) number of 561-Eos (cas9 + *il10rb* gRNA + H2O vs cas9 + *il10rb* gRNA + L-SOP p = 0.7942; unpaired t-test) microglia and (B) number of cannibal microglia (cas9 + *il10rb* gRNA + H2O vs cas9 + *il10rb* gRNA + L-SOP p = 0.2462; unpaired t-test) in 5 dpf cas9 + *il10rb* gRNA animals bath treated with H_2_O or the phagocytosis inhibitor L-SOP. (C) Representative confocal z-projections of 5 dpf *Tg(pu1:Eos)* cas9 + *il10rb* gRNA animals bath treated with H_2_O or L-SOP showing photoconverted microglia and photoconverted debris/presence of cannibal microglia. (D-E) Quantification of the (D) number of 561-Eos microglia (rhIL10 + H2O vs rhIL10 + L-SOP p < 0.0001; unpaired t-test) and (E) number of cannibal microglia (rhIL10 + H2O vs rhIL10 + L-SOP p = 0.0116; unpaired t-test) and in 5 dpf cas9 + *il10rb* gRNA animals bath treated with DMSO or STAT3i. (F) Representative confocal z-projections of 5 dpf *Tg(pu1:Eos)* animals injected with rhIL10 and then bath treated with H_2_O or L-SOP showing photoconverted microglia and photoconverted debris/presence of cannibal microglia. (G-H) Quantification of the average (G) number (Cas9 only vs cas9 + *il10rb* gRNA p = 0.6614; unpaired t-test) and (H) diameter of vacuoles (Cas9 only vs cas9 + *il10rb* gRNA p = 0.2615); unpaired t-test) in cas9 only injected animals and cas9 + *il10rb* gRNA animals. White arrowheads denote 561-Eos+ microglia (C,F). Magenta arrowheads denote 561-Eos+ debris/cannibal microglia (C,F). Scale bars represent 20 μm (C,F).

Conversely, we performed a related experiment to test if the cell death-promoting effect of IL10 over-activation is dependent on phagocytic activity. In this experiment, we again photoconverted six microglia per 4 dpf *Tg(pu1:Eos)* animal. We then injected rhIL10 into the brain ventricles to acutely increase IL10 signaling, and bath-treated the animals with either H2O (control) or the phagocytosis inhibitor L-SOP. As expected, rhIL10 administration significantly reduced the survival of photoconverted microglia in the rhIL10 + H2O group, but the addition of L-SOP rescued the dying microglia. (2.3 ± 1.0 561-Eos microglia per rhIL10 + H2O treated animal; 6.1 ± 0.7 561-Eos microglia per rhIL10 + LSOP treated animal) (Fig 6D). We also saw a similar trend when we quantified the number of cannibal microglia per treatment group, with the L-SOP treated animals having significantly fewer cannibalistic microglia (2.6 ± 1.1 cannibalistic microglia per rhIL10 + H2O treated animal; 1.0 ± 0.8 cannibalistic microglia per rhIL10 + L-SOP treated animal) (Fig 6 E-F). This lack of an additive effect in the *il10rb* gRNA, coupled with the rescue of IL10-induced death by L-SOP, suggests that IL10 signaling acts downstream of, or in parallel with, the initial phagocytic engulfment signal, and that IL10RB and phagocytosis are components of the same cell biological process that regulates the final cell death decision.

Given that the sheer volume of phagocytosed material represents an acute degradative burden that requires active regulation of lysosomal capacity, we first considered that IL10 signaling might be involved in regulating lysosomal dynamics. Phagocytosed material is processed within phagolysosomes, which appear as prominent debris-containing vacuoles in *Tg(pu1:Eos)* animals^38^. Given that these large vacuoles accumulate in the setting of intense phagocytic demand at 4 dpf and previous work confirms they contain various types of neural debris, we assessed them as the likely functional phagolysosomal compartments. However, lacking a specific secondary marker to assign them as phagolysosomes, phagosomes, or autophagic structures, we hereby refer to them simply as vacuoles. We therefore hypothesized that loss of IL10 signaling would alter the number or size of these vacuoles. To test this, we quantified the number of microglial vacuoles in Cas9-only control and *il10rb* gRNA in 4 dpf *Tg(pu1:GAL4;UAS:GFP*) animals, focusing on the number and diameter of vacuoles. This analysis revealed no significant difference in either the number (3.5 ± 2.7 vacuoles per cas9 only animal; 3.2 ± 2.8 vacuoles per *il10rb* gRNA) or the average diameter of vacuoles (6.1 ± 2.8 micrometers per cas9 only microglia; 5.4 ± 2.6 micrometers per *il10rb* gRNA microglia) between the two groups (Fig 6G-H) and thereby inconsistent with the hypothesis that IL10 signaling is required for the physical formation or maturation of the phagolysosome. Based on the morphological consistency between groups, we infer that the IL10 signaling does not regulate the structure of the phagolysosome, which prompted us to next investigate if it plays a role in functional capacity.

Given no gross changes in the number or size of vacuoles, we considered the hypothesis that the biochemistry of the phagocytic machinery could be regulated by IL10. Specifically, given that STAT3 (the key downstream mediator of IL10) is known to regulate v-ATPase pumps^52^, we focused on lysosomal acidification as the most likely functional target. Lysosome pH is the critical determinant of degradative efficiency and has been shown to be a major checkpoint for cell fate^25,68^. We hypothesized that *il10rb* knockdown would impair lysosomal acidification. To explore that, we used LysoTracker Red, a fluorescent probe that specifically accumulates in and labels acidic compartments, to assess lysosomal pH in microglia. We quantified the number of LysoTracker^+^ puncta per microglia in Cas9 + *il10rb* gRNA injected animals at 4 dpf and compared them to Cas9-only control animals. Our analysis revealed that microglia in *il10rb* perturbed animals had significantly fewer LysoTracker^+^ puncta, demonstrating a clear defect in lysosomal acidification when IL10 signaling is disrupted (8.1 ± 3.3 LysoTracker^+^ puncta per cas9 only animal; 4.9 ± 1.9 3 LysoTracker^+^ puncta per Cas9 + *il10rb* gRNA animal) (Fig 7A-B). This key finding establishes a mechanistic link between the IL10RB/STAT3 signaling axis and the failure of lysosomal processing.

**Figure 7.**
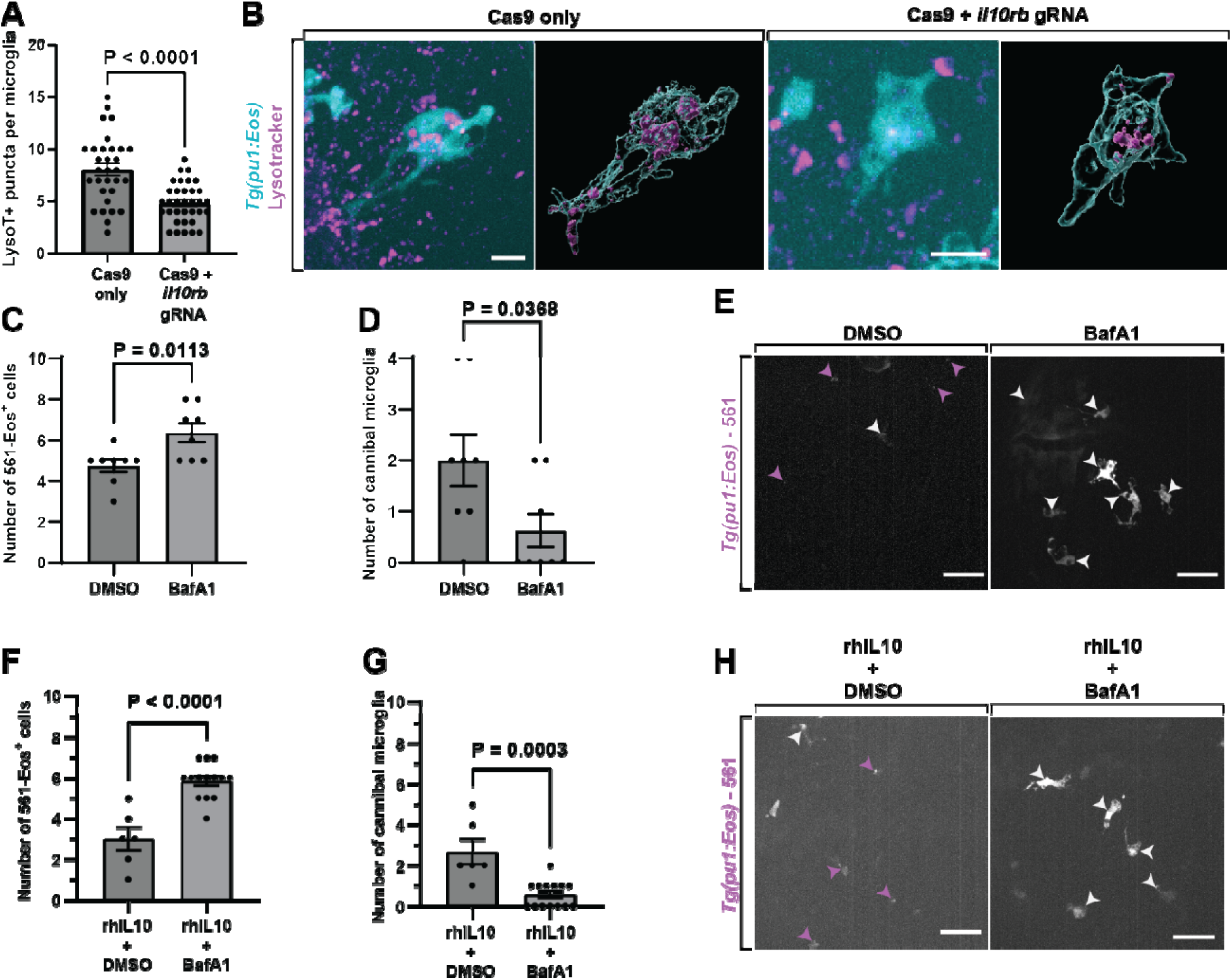
IL-10 Signaling Regulates Microglial Survival via Lysosomal Acidification. (A) Quantification of the number of LysoTracker+ inclusions per microglia (Cas9 only vs cas9 + *il10rb* gRNA p < 0.0001; unpaired t-test) in cas9 only injected animals and cas9 + *il10rb* gRNA animals. (B) Representative confocal z-projections and IMARIS surface renderings of 4 dpf *Tg(pu1:Eos)* animals injected with cas9 only or cas9 + *il10rb* gRNA animals stained with LysoTracker showing acidified lysosome compartments within microglia. (C-D) Quantification of the (C) number of 561-Eos microglia (DMSO vs BafA1 p = 0.0113; unpaired t-test) and (D) number of cannibal microglia (DMSO vs BafA1 p = 0.0368; unpaired t-test) in 5 dpf animals bath treated with DMSO or lysosomal acidification inhibitor BafA1. (E) Representative confocal z-projections of 5 dpf *Tg(pu1:Eos)* animals bath treated with DMSO or BafA1 showing photoconverted microglia and photoconverted debris/presence of cannibalisti microglia. (F-G) Quantification of the number of (F) 561-Eos microglia (rhIL10 + DMSO vs rhIL10 + BafA1 p < 0.0001; unpaired t-test) and (G) cannibalistic microglia (rhIL10 + DMSO vs rhIL10 + BafA1 p = 0.0003; unpaired t-test) in 5 dpf rhIL10 injected animals bath treated with DMSO or BafA1. (H) Representative confocal z-projections of 5 dpf *Tg(pu1:Eos)* animals injected with rhIL10 and bath treated with DMSO or BafA1 showing photoconverted microglia and photoconverted debris/presence of cannibal microglia. White arrowheads denote 561-Eos+ microglia (E,H). Magenta arrowheads denote 561-Eos+ debris/ cannibalistic microglia (E,H). Scae bars represent 10 μm (B) and 20 μm (E,H).

If the acidification of lysosomes is critical for microglia death, potentially regulated via IL10 signaling, then independently altering lysosomal acidification should impact microglial death. To further test the link between lysosomal acidification and microglial death, we used bafilomycin A1 (BafA1), a specific inhibitor of v-ATPase, the proton pump responsible for acidifying lysosomes. We hypothesized that inhibiting acidification would phenocopy the survival phenotype of *il10rb* perturbation. We photoconverted six microglia in 4 dpf *Tg(pu1:Eos)* animals and treated them with either DMSO or BafA1. Consistent with our hypothesis, BafA1-treated animals showed a significant increase in the number of surviving 561-Eos+ cells (4.8 ± 0.9 561-Eos microglia per DMSO treated animal; 6.4 ± 1.3 561-Eos microglia per BafA1 treated animal) and a concomitant decrease in cannibalistic microglia compared to DMSO controls (2.0 ± 1.4 cannibalistic microglia per DMSO treated animal; 0.6 ± 0.9 cannibalistic microglia per BafA1 treated animal) (Fig 7C-E). This provides strong evidence that lysosomal acidification is a key determinant of microglial fate following phagocytosis.

Given that our model predicts that the pro-necroptotic effect of IL10 is mediated by driving lysosomal acidification, we hypothesiszed that pharmacologically blocking acidification would rescue microglia from IL10 induced death. To test this, we injected rhIL10 into the brain ventricles and simultaneously treated the animals with BafA1. The BafA1 treatment successfully negated the pro-death effect of IL10, with survival rates similar to those of BafA1 treatment alone (3.0 ± 1.4 561-Eos microglia per rhIL10 + DMSO treated animal; 5.9 ± 0.9 561-Eos microglia per rhIL10 + BafA1 treated animal), and similarly there were fewer cannibalistic microglia in BafA1 treated animals compared to DMSO treatment (2.7 ± 1.5 cannibal microglia per rhIL10 + DMSO treated animal; 0.6 ± 0.6 cannibal microglia per rhIL10 + BafA1 treated animal) (Fig 7F-H). This is consistent with the hypothesis that the lysosomal acidification pathway is a crucial downstream effector of IL10 signaling in driving microglial death. The simplest explanation for this collective data is that IL10 signaling promotes microglial necroptosis by regulating lysosomal acidification, revealing that the very process required for efficient phagocytosis also serves as the critical checkpoint for triggering cell turnover.

Our findings suggest that IL10 signaling promotes the turnover of phagocytically-burdened cells, establishing a key quality control mechanism. Failure of this mechanism results in the persistence of cells that, though living, are potentially functionally compromised due to the chronic phagocytic stress. Therefore, we next explored the consequences of having longer-lived, *il10rb*-deficient microglia in the context of a neuroimmune challenge. To specifically assess the impact of less microglia turnover on acute injury resolution, we utilized a laser ablation model to induce a focal brain injury in the midbrain, creating a controlled, high-volume source of neural debris that resident microglia must clear. This model was chosen because it allows for a precise, repeatable injury *in vivo*, enabling us to track the dynamics of debris clearance and injury resolution over time, providing a relevant functional readout for the clearance efficiency of the microglial population.

We performed this experiment on 4 dpf *Tg(pu1:Eos)* animals. We photoconverted six microglia per animal, performed a laser ablation injury 50 μm below the surface of the skin in the midbrain, and then re-imaged the same animals at 5 dpf to assess survival and morphology. We compared three groups: wild-type (WT) animals with injury, *il10rb* gRNA-injected animals with injury, and *il10rb* perturbations with a mock injury.

We first examined the overall microglial population in the lesion site to assess the general neuroimmune response to injury. Previous work has shown that the microglia population expands in response to injury. To investigate if *il10rb* perturbation had an effect on this injury response, we quantified the total number of microglia within the imaging window 24 hours post injury (hpi). The total number of microglia in both injured groups (*il10rb* gRNA and WT) was significantly higher than in the mock-injured group, confirming a typical expansion of the microglial population in response to injury, however, there was no difference in the total microglial count between the two injured groups (17.0 ± 5.1 microglia per mock injured *il10rb* gRNA; 25.6 ± 4.9 microglia per injured *il10rb* gRNA; 25.2 ± 3.6 microglia per injured WT animal) (Fig 8A). We had initially hypothesized that because *il10rb* mediates microglia death, *il10rb* perturbed animals would retain a higher number of surviving photoconverted 561-Eos cells. Interestingly, while the mock-injured *il10rb* perturbation retained a high number of 561-Eos microglia (consistent with inhibited microglial developmental death), the injured *il10rb* gRNA-injected animals and WT animals both showed a significant reduction in the number of surviving 561-Eos cells (5.9 ± 0.7 561-Eos microglia per mock injured *il10rb* gRNA; 4.0 ± 1.0 561-Eos microglia per injured *il10rb* gRNA; 3.7 ± 0.8 photoconverted microglia per injured WT animal) (Fig 8B-C). This suggests that the IL10 mediated quality control mechanism that governs developmental microglia turnover is superseded by a more potent, injury-induced cell death pathway, potentially as phagocytic stress reaches a severe acute threshold.

**Figure 8.**
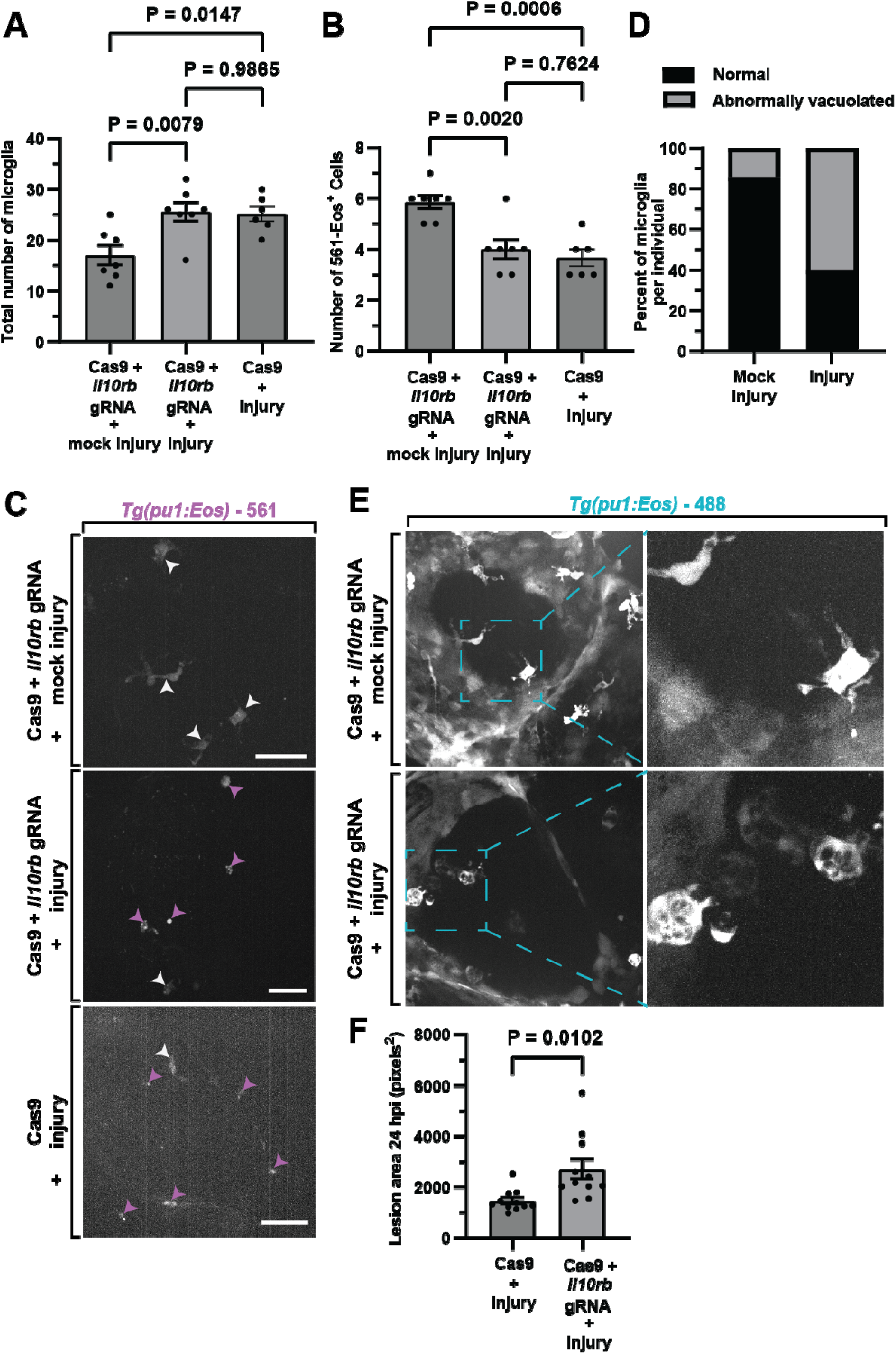
*il10rb* Perturbation Leads to Microglial Dysfunction and Impaired Resolution After Injury. (A-B) Quantification of the (A) total number of microglia (Cas9 + *il10rb* gRNA + mock injury vs. *il10rb* gRNA + injury p = 0.0079; Cas9 + *il10rb* gRNA + mock injury vs. WT + injury p = 0.0147; Cas9 + *il10rb* gRNA + injury vs. WT + injury p = 0.9865; Tukey’s multiple comparisons test) and (B) number of 561-Eos microglia (Cas9 + *il10rb* gRNA + mock injury vs. *il10rb* gRNA + injury p = 0.002; Cas9 + *il10rb* gRNA + mock injury vs. WT + injury p = 0.0006; Cas9 + *il10rb* gRNA + injury vs. WT + injury p = 0.7624; Tukey’s multiple comparisons test) in mock-injured Cas9 + *il10rb* gRNA animals, injured Cas9 + *il10rb* gRNA animals, and injured Cas9 only injected animals. (C) Representative confocal z-projections of 5 dpf *Tg(pu1:Eos)* mock-injured Cas9 + *il10rb* gRNA animals, injured Cas9 + *il10rb* gRNA animals, and injured cas9 only injected animals (D) Quantification of the average percent of normal and highly vacuolated microglia in Cas9 + *il10rb* gRNA animals that were mock-injured or injured. (E) Representative confocal z-projections of 5 dpf *Tg(pu1:Eos)* mock-injured Cas9 + *il10rb* gRNA animals and injured Cas9 + *il10rb* gRNA animals showing microglia morphology. (F) Quantification of the *nbt*- lesion size in 5 dpf *Tg(nbt:dsRed)* Cas9 only injected animals and *il10rb* gRNA injected animals (Cas9 only vs cas9 + *il10rb* gRNA p = 0.0102; unpaired t-test with Welch’s correction). White arrowheads denote 561-Eos+ microglia (C). Magenta arrowheads denote 561-Eos^+^ debris containing cannibal microglia (C). Scale bars represent 20 μm (C).

Next, we investigated whether the failure of IL10 mediated death led to a defect in the remaining microglial function. While we observed that acute injury triggers significant microglial death even in *il10rb* perturbed animals, the surviving cells in these animals that escaped death may have accumulated chronic phagocytic stress from injury clearance. We therefore hypothesized that these surviving, compromised cells would display signs of morphological breakdown and functional failure when exposed to the severe stress of an acute injury. Indeed, we observed a profound difference in microglial morphology following 24 hpi via laser ablation. In the injured *il10rb* perturbed animals, 60% exhibited a highly rounded, vacuolated phenotype while only 40% of microglia displayed a normal, ramified morphology (Fig. 8D-E). In contrast, microglia in the mock-injured *il10rb* gRNA maintained a largely normal appearance (85.7% showing normal morphology). Crucially, we found virtually no microglia with this abnormally vacuolated morphology in the injured WT control group (Fig. 8D-E). This direct comparison highlights that the vacuolated phenotype is not a generic response to injury but is specifically linked to the absence of proper IL10 signaling. This observation strongly suggests that the vacuolated phenotype represents a state of chronic cellular stress and dysfunctional debris clearance in the long-lived, *il10rb*-deficient cells.

To evaluate the consequence of this dysfunctional microglial state, we quantified the resolution of the tissue lesion following injury. We performed the same focal laser ablation in *Tg(nbt:dsRed)* animals, which fluorescently labels neurons, allowing us to quantify the size of the tissue lesion. We compared Cas9-only controls to *il10rb* gRNA 24 hours post-injury. The lesion size was significantly larger in *il10rb* gRNA compared to Cas9-only controls (1452 ± 435.3 pixels^2^ per cas9 only animal; 2703 ± 1297 pixels^2^ per *il10rb* gRNA) (Fig. 8F). This demonstrates that the failure to execute the IL10RB-driven quality control mechanism—resulting in the persistence of longer-lived, but dysfunctional, microglia—directly leads to impaired debris clearance and delayed resolution of tissue damage *in vivo*.

## Discussion

This work supports a model that IL10 signaling promotes microglial death. This process is essential for maintaining a healthy, functional population of microglia capable of responding effectively to injury and clearing debris. The longer-lived microglia when IL10 signaling is disrupted are functionally compromised under conditions of high phagocytic demand, exhibiting characteristics of resolution failure due to impaired lysosomal function. This further supports the model that the IL10/necroptosis paradigm is a quality control mechanism designed to eliminate stressed, defective phagocytes to ensure optimal tissue repair and remodeling.

Overall, we hypothesize that during periods of intense phagocytic demand in early brain development, IL10 signaling functions as a direct mechanism to refine the microglia population of phagocytically burdened cells. Following IL10 binding, JAK/STAT3 signaling is initiated. Although not directly linked to IL10 as the signal initiator, work has found that activated STAT3 translocates to the phagolysosome membrane and physically associates with the V-ATPASE complex, directly promoting rapid acidification^52^. While initially protective and helpful for cargo degradation, we suspect the chronic demand leads to the hyper-activation of this STAT3-mediated acidification. Mechanistically, we propose this sustained hyper-acidification, combined with the accumulation of debris, could compromise lysosomal membrane integrity, triggering LMP, which is a known driver of necroptosis^69^. This catastrophic failure releases lysosomal contents into the cytosol, which could initiate the observed necroptotic cell death pathway in microglia^38^. Our finding that *il10rb* knockdown reduces microglial death is therefore consistent with removing the signal that actively drives the lysosomal system towards necroptosis.

### IL10 Signaling Governs Microglial Turnover via a Lysosomal Quality Control Mechanism

In this study, we utilized a high-throughput CRISPR-Cas9 screen and single-cell *in vivo* tracking in the larval zebrafish to identify the molecular mechanisms regulating microglial turnover during development. Our key findings establish a signaling cascade wherein the cytokine IL10 acts as a pro-death signal through its accessory receptor, IL10RB, to actively promote the necroptosis of phagocytic microglia. Mechanistically, we demonstrate that this effect is mediated through the canonical JAK/STAT3 axis, and that IL10 signaling is essential for maintaining lysosomal acidification. Building on previous literature that establishes that STAT3 physically associates with and regulates the v-ATPase proton pump^52^, we hypothesize that this is the direct molecular link by which the axis control lysosomal pH. By tuning this critical pH checkpoint, the IL10-IL10RB-JAK-STAT3 axis links the stress of intense efferocytosis to a regulated cell death decision, resulting in the cannibalistic clearance of necroptotic microglia. We propose that this IL10-driven necroptosis process functions as a self-limiting quality control mechanism to ensure the developing brain maintains a population of healthy, functional phagocytes.

### IL10-STAT3 Axis as a Core Regulator of Lysosomal Integrity and Cell Fate

Our work provides a crucial mechanistic link between a major inflammatory signaling pathway and the ultimate decision of cell death. The convergence of IL10/STAT3 signaling on lysosomal function is particularly compelling. Canonical STAT3 activity is generally associated with cell survival and proliferation^70,71^, yet we found that inhibiting STAT3 rescued microglial lifespan (Fig. 5). This seemingly contradictory role is resolved by our discovery that the IL10-IL10RB-JAK-STAT3 signaling axis is required to maintain the proper acidic environment (pH) of the lysosome (Fig. 7). This finding strongly supports a mechanism, established in previous literature, which shows that STAT3 directly associates with the v-ATPase proton pump and increases its activity^52^. In the context of the highly burdened phagocytic microglia, we hypothesize that IL10-mediated acidification is a double-edged sword: it is required for successful debris degradation, but in the face of continuous, high-volume efferocytosis, this intense v-ATPase activity also drives the lysosome toward a state of fatal stress, leading to eventual lysosomal membrane permeabilization (LMP) and the initiation of RIPK1-dependent necroptosis^72,73^. This is strongly supported by our finding that the microglia death effect of exogenous IL10 is abrogated by Nec−1 treatment, a specific pharmacological inhibitor of RIPK1 (Fig 4). Therefore, IL10 does not promote death directly, but rather tunes the lysosome to a high-activity state that is necessary for function but predisposes the cell to death under overwhelming stress. The demonstration that BafA1 (a v-ATPase inhibitor) completely phenocopied the survival effect of *il10rb* knockdown (Fig. 5), and that it eliminated the microglia death effect of exogenous IL10, strongly supports the lysosomal pH as a critical execution point of this developmental mechanism. The IL10 signal, therefore, acts as a systemic regulator of phagocytic efficiency and risk, essentially setting the threshold for resolution failure.

### Linking Developmental Turnover to Disease Resolution

Our finding that *il10rb*-deficient microglia, while longer-lived, exhibit severe morphological dysfunction and a highly vacuolated phenotype following injury (Fig. 8) has significant implications for our understanding of neuroimmune cell longevity in disease. This phenotype is indicative of a stalled clearance state or phagocytic overload, where the microglia survive the initial death signal but are functionally incapable of resolving their internalized cargo. The failure of phagocytes to die and be replaced is increasingly recognized as a major driver of chronic, non-resolving inflammation in pathologies such as atherosclerosis^26,28,35,37^ and advanced aging^74,75^. Our model suggests that the IL10-driven necroptosis is a developmentally conserved mechanism to prevent the accumulation of these dysfunctional, "resolution-failed" phagocytes. By rapidly eliminating the damaged cells, the genetic pathway ensures the tissue is populated by a new, high-functioning cohort of microglia capable of responding effectively to subsequent challenges. This paradigm moves beyond viewing IL−10 simply as an anti-inflammatory agent, revealing its capacity to orchestrate the strategic removal of immune cells as a prerequisite for tissue-level resolution.

### Limitations and Future Directions

While this work establishes a clear signal-to-fate pathway, several questions remain. Our study focuses on the IL10RB subunit, which pairs with IL10Rα to form the functional receptor; future work should explore the specific role of IL10RA and the full heterodimer complex. Furthermore, although we demonstrated a strong link between IL10 and lysosome acidification, the precise molecular nature of their interaction within microglia will need to be tested in future publications. Future studies should also confirm that STAT3 physically associates with the v-ATPase complex in microglia. Finally, given that IL10 is broadly expressed, it will be crucial to distinguish the contributions of the microglial cell-autonomous IL10 signal from potential signals originating from neurons, glia, or other cell types, especially under conditions of injury. Ultimately, translating this developmental mechanism to adult disease models will determine whether pharmacologically targeting the IL10/STAT3/v-ATPase axis can be used to promote the necessary turnover and rejuvenation of phagocyte populations in chronic inflammatory conditions.

## Author Contribution

HG, JD, AD, ZMK, RAA, KC, CC, and CH performed experimentation.

HG and DG performed the analysis.

HG, JD, AD, and CJS supervised the project.

HG and CJS conceived, wrote, and edited the manuscript and funded the project.

## Acknowledgements

We thank Beth Stevens, Jordan Doman, Anna Kane and members of the Stevens’ lab for their thoughtful comments. We also thank current and previous members of the Smith Lab for insightful discussions. Thanks to 3i for aiding with imaging related questions, Sara Cole of the Notre Dame Integrated Imaging Facility for assisting with IMARIS, and Deborah Bang for zebrafish housing and upkeep. This work was supported by the University of Notre Dame, the Elizabeth and Michael Gallagher Family, Centers for Zebrafish Research and Stem Cells and Regenerative Medicine at the University of Notre Dame, the SMART foundation (CJS), NIH (DP2NS117177) and Chan Zuckerberg Initiative (CP-2-1-Smith).

---

**Figure S1.**
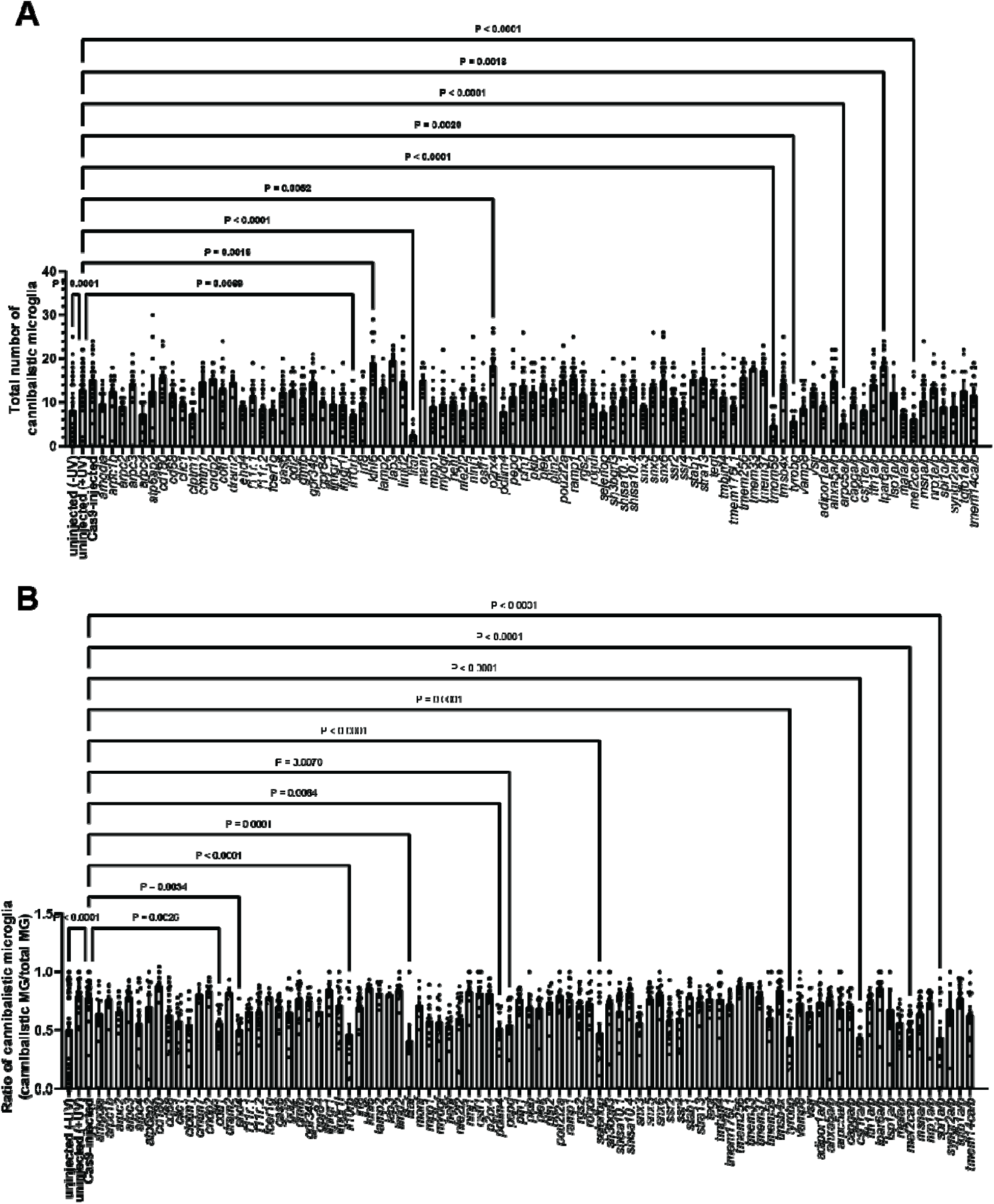
Large scale CRISPR-Cas9 screen reveals perturbations in microglia cannibalism upon gene knockdown. (A-B) Quantification of (A) the number of cannibalistic microglia (Cas9-injected vs. *uninjected* (-UV) p < 0.0001; Cas9-injected vs. *arpc2* p = 0.0089; Cas9-injected vs. *arpc4* p = 0.0003; Cas9-injected vs. *clic1* p = 0.0227; Cas9-injected vs. *clptm1* p < 0.0001; Cas9-injected vs. *ehd4* p = 0.0159; Cas9-injected vs. *f11r.2* p < 0.0001; Cas9-injected vs. *gmfb* p = 0.0376; Cas9-injected vs. *ifngr1l* p = 0.0009; Cas9-injected vs. *il10rb* p < 0.0001; Cas9-injected vs. *irf8* p = 0.0038; Cas9-injected vs. *litaf* p < 0.0001; Cas9-injected vs. *mpp1* p = 0.0072; Cas9-injected vs. *mydgf* p = 0.01; Cas9-injected vs. *nfe2l2* p < 0.0001; Cas9-injected vs. *ostf1* p = 0.0237; Cas9-injected vs. *pdlim4* p = 0.0011; Cas9-injected vs. *rogdi* p = 0.015; Cas9-injected vs. *selenop* p < 0.0001; Cas9-injected vs. *sh3bgrl3* p = 0.0014; Cas9-injected vs. *shisa10.1* p = 0.0436; Cas9-injected vs. *snx3* p = 0.0024; Cas9-injected vs. *ssr4* p = 0.0002; Cas9-injected vs. *tmem176l.1* p < 0.0001; Cas9-injected vs. *tmem59* p < 0.0001; Cas9-injected vs. *tyrobp* p < 0.0001; Cas9-injected vs. *vamp8* p = 0.0049; Cas9-injected vs. *adipor1a/b* p = 0.0005; Cas9-injected vs. *arpc5a/b* p < 0.0001; Cas9-injected vs. *capga/b* p = 0.0044; Cas9-injected vs. *csf1ra/b* p = 0.0038; Cas9-injected vs. *mafa/b* p = 0.0001; Cas9-injected vs. *mef2ca/b* p < 0.0001; Cas9-injected vs. *msna/b* p = 0.0172; Cas9-injected vs. *spi1a/b* p = 0.0004; Dunnett’s multiple comparisons test) and (G) the ratio of cannibalistic microglia (Cas9-injected vs. *uninjected* (-UV) p < 0.0001; Cas9-injected vs. *clic1* p = 0.043; Cas9-injected vs. *clptm1* p = 0.014; Cas9-injected vs. *cotl1* p = 0.0026; Cas9-injected vs. *ehd4* p = 0.0034; Cas9-injected vs. *il10rb* p < 0.0001; Cas9-injected vs. *litaf* p = 0.0001; Cas9-injected vs. *mydgf* p = 0.0196; Cas9-injected vs. *nenf* p = 0.0223; Cas9-injected vs. *pdlim4* p = 0.0064; Cas9-injected vs. *pepd* p = 0.007; Cas9-injected vs. *selenop* p < 0.0001; Cas9-injected vs. *snx3* p = 0.0259; Cas9-injected vs. *ssr2* p = 0.018; Cas9-injected vs. *tyrobp* p = 0.0001; Cas9-injected vs. *csf1ra/b* p < 0.0001; Cas9-injected vs. *mef2ca/b* p < 0.0001; Cas9-injected vs. *spi1a/b* p < 0.0001; Dunnett’s multiple comparisons) compared to the total microglia population upon gene knockdown with CRISPR-Cas9 in 4 dpf *Tg(pu1:Eos)* animals.

**Figure S2.**
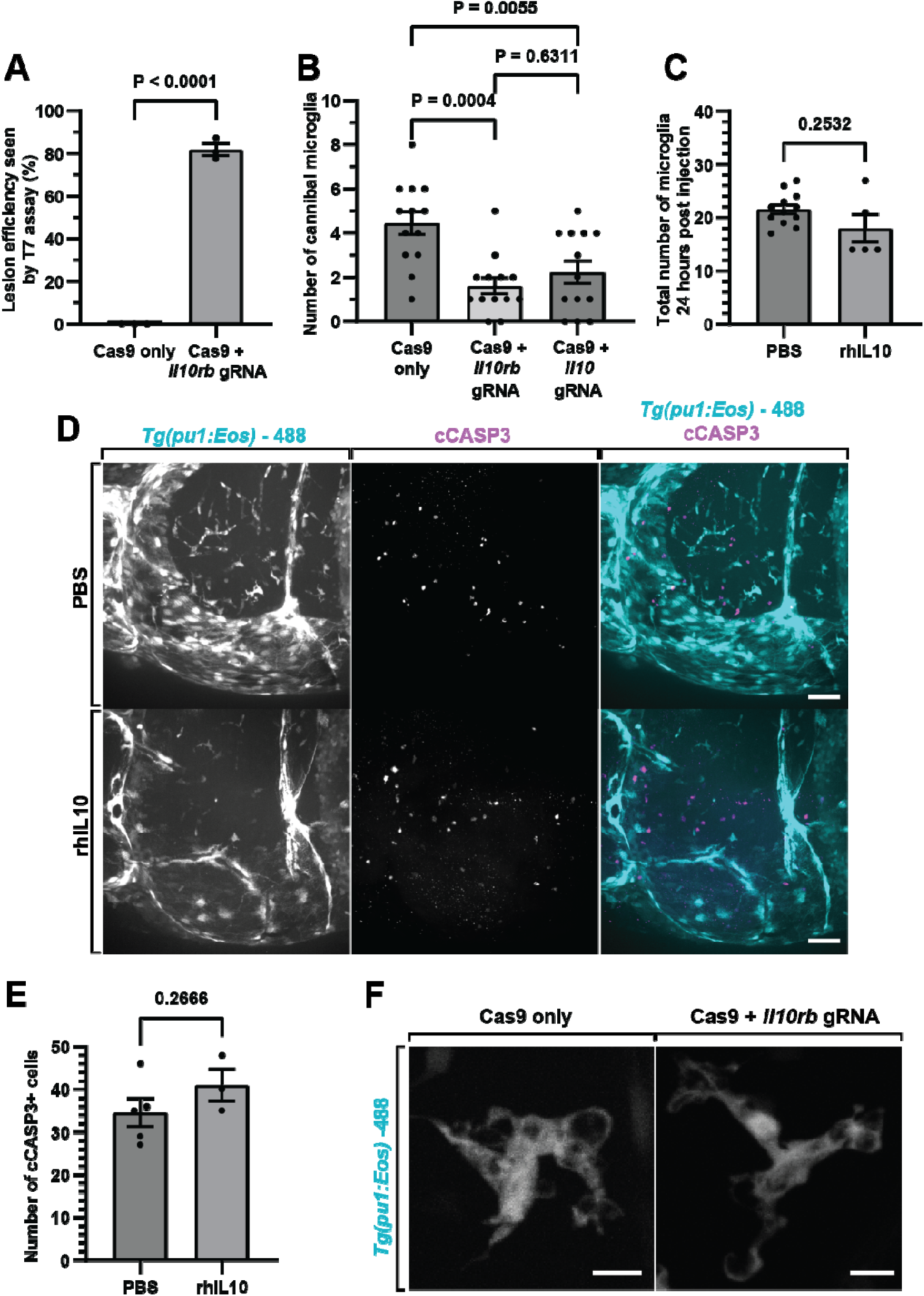
*il10rb* + *il10* Perturbation Leads to Microglial Dysfunction. (A) Quantification of CRISPR-Cas9 lesion efficiency for *il10rb* gRNA (Cas9 only vs Cas9 + *il10rb* gRNA p < 0.0001; unpaired t-test). (B) Quantification of the number of cannibalistic microglia in Cas9 only, Cas9 + *il10rb* gRNA, or Cas9 + *il10* gRNA (Cas9 only vs. Cas9 + *il10rb* gRNA p = 0.0004, Cas9 only vs. Cas9 + *il10* gRNA p = 0.0055, Cas9 + *il10rb* gRNA vs. Cas9 + *il10* gRNA p = 0.6311, Tukey’s multiple comparisons test). (C) Quantification of the number of microglia PBS and rhIL10 injected animals 24 hours post injection (PBS vs rhIL10 p = 0.2532; unpaired t-test). (D) Representative confocal z-projections of 5 dpf *Tg(pu1:Eos)* animals injected with PBS or rhIL10 showing cCASP^+^ cells and microglia. (E) Quantification of the total number of cCASP^+^ cells in 5 dpf *Tg(pu1:Eos)* animals injected with PBS or rhIL10 (PBS vs rhIL10 p = 0.2666; unpaired t-test). (F) Representative confocal z-projections of 5 dpf *Tg(pu1:Eos)* animals injected with cas9 only or cas9 + *il10rb* gRNA showing similar vacuole morphology.

## Methods

### Ethics Statement

Experimental procedures followed the NIH guide for the care and use of laboratory animals. The University of Notre Dame IACUC, which is guided by the United States Department of Agriculture, the Animal Welfare Fact (USA), and the Assessment and Accreditation of Laboratory Animal Care International approved all animal studies under protocol 22-07-7322 and 25-05-9326 to Cody J. Smith.

### Contact for Reagent and Resource Sharing

All data collected for the study are included in the figures and supplemental data. Reagent requests can be directed to the corresponding author.

### Custom Code

Custom code used to generate the figures can be found at https://github.com/RavenGan/Microglia.

### Experimental Model and Subject Details

Zebrafish strains in this study include: AB (wild-type), *Tg(pu1:Eos)^nt20076^*, *Tg(pu1:GAL4, UAS:GFP)^zf1496^*, and *Tg(nbt:dsRed)^zf1486^*. Only stable transgenic lines were used in this study. To produce embryos, pairwise matings were used. Animals were raised at 28 °C in egg water in constant darkness and staged by hours or days post fertilization (hpf and dpf), confirmed by observation of developmental milestones^77^. Embryos were raised in 0.003% phenylthiourea (PTU) egg water from 24 hpf – 4 dpf to prevent pigment development. Embryos were used for all experiments.

### Method Details

#### scRNAseq Analysis

The microglia scRNAseq data were downloaded from GSE121654^32^. Pre-processing, normalization, and clustering were performed using Seurat as described previously^78^.

#### In Vivo Imaging

Animals were anesthetized using 3-aminobenzoic acid ester (Tricaine), and submerged in 0.8% low-melting point agarose, and mounted dorsally in glass-bottomed 35 mm Petri dishes as previously described^79^. Custom confocal microscopes, built by 3i technology (Denver, Colorado, United States of America) was used for all imaging. All mounted animals were imaged with a 40X (1.1NA) objective. Images were processed with Adobe Illustrator, ImageJ, and IMARIS. Only brightness and contrast were adjusted and enhanced for images represented in this study.

### Screening Robot Imaging

A robot equipped with a Vast BIOImager (Union Biometrica) and a custom built confocal microscope from 3i technology (Denver, Colorado) was used for high-throughput screening^78^. The microscope includes: Zeiss Axio Examiner Z1, CSU-W1 T1 Spinning Disk Confocal (50 μm disk) equipped with mSwitcher, dichroics and lasers for 405/488/561/640 and a 95B Back Illuminated Scientific CMOS. Animals were anesthetized and placed in a 96-well plate prior to imaging.

### Immunohistochemistry and Hybridization Chain Reaction (HCR)-Fluorescence In Situ Hybridization (FISH)

The primary antibodies used were anti-4C4 (1:50, mouse^65^) and anti-cleaved Caspase3 (1:700, rabbit, BD Biosciences). The secondary antibodies used were Alexa Fluor 647 goat anti-mouse (1:600, Thermo Fisher, A-21235) and Alexa Fluor 647 goat anti-rabbit (1:600, Thermo Fisher, A-21245). Animals were fixed at 4 dpf and 5 dpf in 4% paraformaldehyde in 0.1% PBS Triton-X. HCR-FISH was used in combination with IHC and was performed using the standard protocol (Molecular Instruments, HCR RNA-FISH v3.0 protocol for whole-mount zebrafish larvae) on 4 dpf fixed animals to detect *irf8*, *clic1*, *nfe2l2*, *arpc4*, *mef2ca/b*, *il10*, and *il10rb* expression. Probes targeting these genes were custom designed.

### CRISPR-Cas9 Screen and Knockdown

#### Targeted CRISPR-Cas9 Mutagenesis for Screen

CRISPR-Cas9 mutagenesis was performed by co-injecting a pool of 3-4 guide RNAs (gRNAs) targeting a single gene and 2 μM EnGen Spy Cas9 NLS (New England Biolabs) into *Tg(pu1:Eos)* embryos at the 1-cell stage. Candidate genes for knockdown were chosen from a re-analysis of a published scRNA-seq dataset of E14.5 mouse microglia^41^. gRNAs were made using the online program CHOPCHOP. The following parameters were used: gRNAs are be 20 bp in length, begin with GA, and have no off targets with fewer than 3 bp matches.

#### Screen Workflow

After gRNA and Cas9 injection, injected embryos were grown to 12 hours post fertilization (hpf). To allow for mosaic labeling of microglia, at 12 hpf, embryos were exposed to 405 nm light using two, 5 second pulses of light from an LED flood system (Loctite 97070). After photoconversion, animals were grown to 4 dpf in PTU egg water. The 4 dpf animals were randomly selected and imaged on a custom-built robot. Hits were validated using the same CRISPR-Cas9 protocol with individual gRNA pools.

#### Targeted CRISPR-Cas9 Mutagenesis for *il10* and *il10rb*

CRISPR-Cas9 mutagenesis was performed by co-injecting two guide RNAs (gRNAs) targeting a single exon within each gene and Alt-R S.p. Cas9 Nuclease V3 protein (Integrated DNA Technologies, 1081058) into *Tg(pu1:Eos)* embryos at the 1-cell stage. Crispants were confirmed with T7 endonuclease assay.

#### T7 Endonuclease Assay

A T7 endonuclease I (New England Biolabs, M0302SVIAL) assay was performed to determine if an indel was created at or near the exon of interest and to determine guide efficiency. The assay was carried out according to New England Biolabs protocol instructions. Only animals with digested PCR product (indicating an indel) were kept in for analysis.

#### Six-cell Photoconversions and Survival Tracking

*Tg(pu1:Eos)* embryos at 4 dpf were used. Six individual microglia in the midbrain were selected and photoconverted using a 5-ms 405-nm-laser pulse with the mVector system (3i). Pre- and post-conversion images were taken in the 488 nm and 561 nm channels. Survival was assessed by tracking the persistence of the 561-Eos cells and by the presence of cannibal microglia at 5 dpf.

### Drug Treatments

#### Chemical Bath Treatments

Working solutions were administered by diluting the drug in PTU egg water at 4 dpf immediately after photoconversion. Signaling Inhibitors: A pan JAK inhibitor (JAKi; CAS 457081-03-7; Sigma-Aldrich, Cat# CC1000) was used at 1 μM^80^. A STAT3 inhibitor (STAT3i; S31-201; Sigma-Aldrich, Cat# 573130) was used at 400 μM^80^. Cell Death Inhibitors: Necrostatin-1 (Nec-1; RIPK1 inhibitor; MedChemExpress LLC, HY-15760) was used at 10 μM^38^. Lysosome acidification inhibitor: the selective inhibitor of the vacuolar-type H^+^-ATPase, Bafilomycin A1 (BafA1;Sigma-Aldrich, SML1661) was used at 25 nM^81^. Phagocytosis Inhibitor: O-Phospho-L-serine (L-SOP; Sigma, Sigma, P0878-10MG) was used at 1 μM^38^. All embryos were incubated in egg water until 24 hpf and incubated with PTU until desired treatment time. Fish were treated at 4 dpf, immediately after photoconversions and/or injections. Control fish were incubated with 0.1% DMSO in PTU or H_2_O in PTU.

#### rhIL10 Injection

Following the photoconversion protocol, embryos were carefully unmounted from their dorsal position. They were then briefly re-anesthetized using Tricaine in PTU egg water and remounted ventrally (head facing up) in a glass-bottom dish with a small amount of low-melt agarose to stabilize the head and expose the brain ventricle. Recombinant human IL10 (rhIL10; R&D Systems; 217-IL-005/CF) was injected into the brain ventricle at 25 μg/ml using a glass micro-injection needle connected to a micro-injector. Vehicle control injections consisted of PBS. Following injection, embryos were immediately unmounted and placed in PTU egg water or the appropriate bath treatment (e.g., Nec-1, BafA1, etc.) to recover until 5 dpf imaging.

### Lysosomal Acidification and Vacuole Quantification

#### LysoTracker Staining

Lysosomal acidification was assessed by bathing 4 dpf animals in LysoTracker Red DND-99 (Thermo Fisher; L7528) at 1 μM for 1 hour prior to imaging. Embryos were rinsed three times with PTU egg water prior to mounting. LysoTracker-positive puncta within individual microglia were manually counted across the z-stack.

#### Vacuole Quantification

Vacuole number and size were quantified in *Tg(pu1:GAL4; UAS:GFP)* fish at 4 dpf. Vacuoles were defined as GFP^-^ inclusions fully contained within the GFP^+^ cytoplasm of the microglia.

#### Laser Ablation Injury

Focal brain injury was induced in *Tg(pu1:Eos)* 4 dpf animals using the Ablate! Photoablation System (532 nm pulsed laser) in the midbrain 40 μm below the skin surface as previously described^38^. Sham-injured fish followed the same mounting procedure but received only a 561 nm laser exposure.

### Quantification and Statistics

#### Statistical Analysis

Microglia z-stack images were generated using 3i Slidebook software. Three-dimensional surface renderings of microglia were created using IMARIS (Notre Dame Imaging Core). All quantifications were executed using various plug-ins available in FIJI (ImageJ) and Microsoft Excel. Statistical analyses were performed using GraphPad Prism (version 8) software. All graphical data represent both the mean and individual values unless otherwise noted.

No statistical methods were used to predetermine sample sizes; sample sizes were informed by previous publications. All statistical tests were run using biological replicates, not technical replicates. Healthy animals were randomly selected for inclusion, and no data points were excluded from analysis. Data distribution was assumed to be normal, but this was not formally tested. Data collection and analysis were performed blind to the experimental conditions unless otherwise indicated. Each experiment was repeated at least twice with similar results.

### Softwares

ImageJ and Slidebook were used to produce and process confocal images. IMARIS was used to generate 3D surface renderings. Graphpad prism was used to generate all graphs and statistical analysis. Adobe Illustrator was used to compile the figures. Gemini was used for editing of the manuscript. All text in the manuscript has been reviewed by the authors for accuracy after Gemini edits.

